# Circuit mechanisms for the maintenance and manipulation of information in working memory

**DOI:** 10.1101/305714

**Authors:** Nicolas Y. Masse, Guangyu R. Yang, H. Francis Song, Xiao-Jing Wang, David J. Freedman

## Abstract

Recently it has been proposed that information in short-term memory may not always be stored in persistent neuronal activity, but can be maintained in “activity-silent” hidden states such as synaptic efficacies endowed with short-term plasticity (STP). However, working memory involves manipulation as well as maintenance of information in the absence of external stimuli. In this work, we investigated working memory representation using recurrent neural network (RNN) models trained to perform several working memory dependent tasks. We found that STP can support the short-term maintenance of information provided that the memory delay period is sufficiently short. However, in tasks that require actively manipulating information, persistent neuronal activity naturally emerges from learning, and the amount of persistent neuronal activity scales with the degree of manipulation required. These results shed insight into the current debate on working memory encoding, and suggest that persistent neural activity can vary markedly between tasks used in different experiments.

## Introduction

Working memory refers to our ability to temporarily maintain and manipulate information, and is a cornerstone of higher intelligence^1^. In order to understand the mechanisms underlying working memory (WM), we must resolve the substrate(s) in which information in working memory is maintained. It has been assumed that information in WM is maintained in persistent neuronal activity^2–6^, likely resulting from local recurrent connections^7,8^, and/or cortical to subcortical loops^9^. However, recent experiments reveal that the strength of persistent neuronal activity varies markedly between tasks^10–16^. This raises two related questions: 1) why does persistent neural activity vary between tasks, and 2) for those tasks with weak or non-existent persistent activity, where and how is information maintained?

A possible answer to the second question is that information is not necessarily maintained in persistent activity, but can be maintained through short-term synaptic plasticity (STP). STP, which is distinct from more commonly known long-term depression (LTD) and potentiation (LTP), is the process in which pre-synaptic activity increases the calcium concentration in the presynaptic terminal but depletes neurotransmitter stores, altering synaptic efficacies on timescales of hundreds of milliseconds to seconds^17^. Importantly, modelling studies have suggested that STP can allow networks to maintain an “activity-silent” memory trace of a stimulus, in which short-term information is maintained without persistent activity^18^. Recent work in human subjects suggests that information can be mnemonically encoded in a silent, or latent, state, and that information can be reactivated into neuronal activity by probing the circuit^19,20^.

While STP might provide another mechanism for the maintenance of information, it does not in itself fully account for why the strength of persistent activity varies between tasks. To answer this, we highlight that WM involves not just the maintenance of information, but also its manipulation. Importantly, manipulating information in short-term memory appears to engage the frontoparietal network differently compared to simply maintaining information^21,22^. While STP can allow for activity-silent maintenance of information, it is unknown whether STP can support activity-silent manipulation of information without persistent activity. If it cannot, then it suggests that the strength of persistent activity reflects the degree of information manipulation required by the task.

In this study, we examine whether STP can support the silent manipulation of information in WM, and whether it could explain the variability in the strength of persistent activity between tasks. Unfortunately, with current technology it would be technically daunting to measure synaptic efficacies in awake behaving mice, and next to impossible in non-human primates. However, recent advances in recurrent neural network (RNN) model algorithms have opened an entirely new avenue to study the putative neural mechanisms underlying various cognitive functions. Crucially, RNN models have successfully reproduced the patterns of neural activity and behavioral output that are observed *in vivo*, and have generated novel insights into neural circuit function that would otherwise be unattainable through direct experimental measurement^23–29^.

Here, we train biologically inspired RNN models, consisting of excitatory and inhibitory like neurons^30^ and dynamic synapses governed by STP^18^, to solve a variety of widely studied WM based tasks. These tasks involved maintaining information (Figure 2), manipulating the contents of short-term memory (Figures 3&4), changing how information is represented (Figure 5), and attending to specific memoranda (Figure 6).

We show that STP can support the activity-silent maintenance of information, but that it cannot support the silent manipulation of information without persistent activity. Furthermore, we show that the strength of persistent neural activity covaries with the degree of manipulation. This potentially explains the widely observed observation that persistent activity varies markedly between tasks.

## Results

The goal of this study was 1) to determine whether STP can support activity-silent manipulation of information in WM (Figures 2–6), and 2) whether STP can explain the variability in persistent activity observed in different tasks^10–16^ (Figure 7). We trained RNN models to solve several widely studied WM tasks, which varied in their specific cognitive demands (see below). In all tasks, the stimuli were represented as coherent motion patterns moving in one of eight possible directions. However, the results of this study are not meant to be specific to motion, or even visual, inputs, and the use of motion patterns as stimuli was simply used to make our example tasks more concrete. Furthermore, similar to the metabolic constraints faced by real brains^31^, we added a penalty on high neuronal activity (see Methods) so that networks were encouraged to solve tasks using low levels of neuronal activity.

### Network model

We constrained neurons in our network to be either excitatory or inhibitory^30^. The input layer of the network consisted of 36 excitatory, direction tuned neurons, whose responses across directions were modelled as a Von Mises function (see Methods). These input neurons projected onto a recurrently connected network of 80 excitatory and 20 inhibitory neurons (Figure 1A). Recurrent neurons never sent projections back onto themselves.

**Figure 1.**
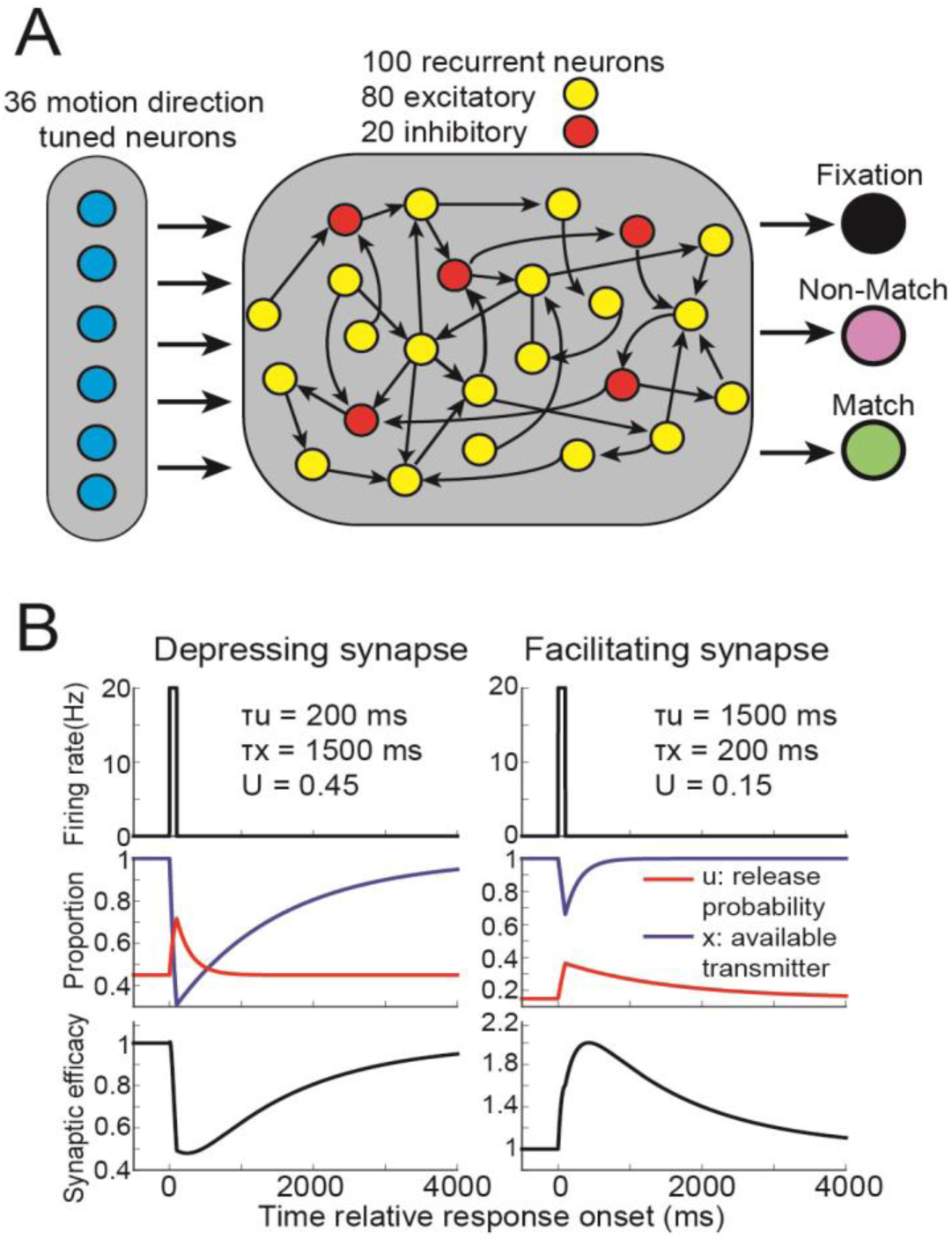
Recurrent neural network design. **(A)** The core rate-based network model consisted of 36 motion direction tuned neurons that sent non-negative projections onto 100 recurrently connected neurons. Of these 100 neurons, 80 were excitatory (their output weights were non-negative) and 40 were inhibitory (their output weights were non-positive). The 80 excitatory neurons sent non-negative projections onto three decisions neurons. **(B)** For synapses that exhibited short-term synaptic depression (left panels), a 100 ms burst of presynaptic activity at 20 Hz (top panel) will increase the value representing the synaptic release probability (red trace, middle panel) and decrease the value representing the available neurotransmitter (blue trace). For these depressing synapses, the stronger and longer lasting decrease in release probability will lead to a decrease in synaptic efficacy (bottom panel) lasting thousands of milliseconds. For synapses that exhibited short-term synaptic depression (right panels), the same burst of presynaptic activity will lead to a stronger and longer lasting increase in release probability (middle panel), leading to an increase in synaptic efficacy lasting thousands of milliseconds.synaptic efficacy lasting thousands of milliseconds (right panel). For computational efficiency, these values will be identical for all synapses sharing the same presynaptic neuron.

The connection weights between all recurrently connected neurons, were dynamically modulated by short-term synaptic plasticity (STP, see Methods) using a previously proposed model^18^. In this model, synaptic efficacy is proportional to the product of two terms: one representing the amount of available transmitter, and another representing the release probability. Connection weights from half of the neurons were depressing, such that presynaptic activity strongly decreases the amount of available neurotransmitter but only weakly increases release probability, leading to a decrease in synaptic efficacy lasting thousands of milliseconds (Figure 1B, left panel). In contrast, connection weights from the other half of the neurons were facilitating, such that presynaptic activity weakly decreases the amount of available neurotransmitter and strongly increases the release probability, leading to an increase in

### Maintaining information in short-term memory

We first examined how networks endowed with STP maintain information in WM using either persistent neuronal activity or using STP. We trained 20 networks to solve the delayed match-to-sample task (DMS, Figure 2A), in which the networks had to indicate whether sequentially presented (500 ms presentation; 1000 ms delay) sample and test stimuli were an exact match. Task accuracy for all networks in this study was >90%.

**Figure 2.**
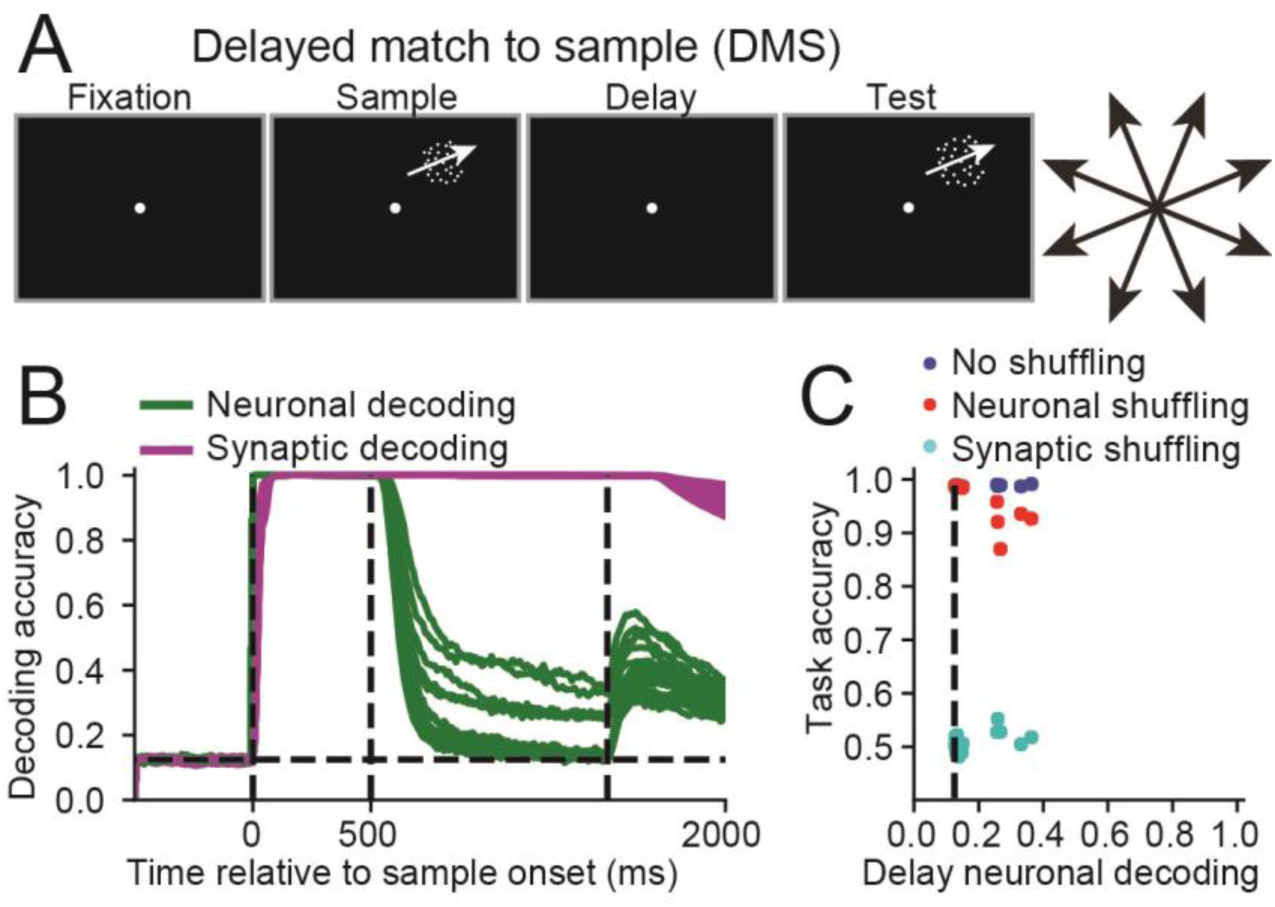
Neuronal and synaptic mnemonic encoding for the delayed match-to-sample task. **(A)** In the delayed match-to-sample (DMS) task, a 500 ms fixation period was followed by a 500 ms sample stimulus, which is represented as a coherent random dot motion pattern which could move in one of eight directions. This was followed by a 1000 ms delay period and finally a 500 ms test stimulus, which was also represented as motion in one of eight directions. The network was trained to indicate whether the motion directions of the sample and test stimuli matched. (B) The time course of the sample direction decoding accuracy, calculated using neuronal activity (green curves) or synaptic efficacy (magenta curves) are shown for all twenty networks that successfully solved the DMS task. The dashed vertical lines, from left to right, indicate the sample onset, offset, and end of the delay period. **(C)** Scatter plot showing the neuronal decoding accuracy measured at the end of the delay (x-axis) versus the behavioral accuracy (y-axis) for all 20 networks trained on this task (blue circles), the behavioral accuracy for the same 20 networks after neuronal activity was shuffled right before test onset (red circles) or synaptic efficacies were shuffled right before test onset (cyan circles). Thus, for each blue circle, there is a corresponding red and cyan circle with matching neuronal decoding accuracy. The dashed vertical line indicates chance level decoding accuracy.

To measure how information was maintained, we linearly decoded the sample direction using support vector machines (SVM) that were trained and tested at 10 ms intervals, from 1) the population activity of the 100 recurrent neurons, and from 2) the 100 unique synaptic efficacy values dynamically modulated by STP. Classifiers were individually trained and tested at each 10 ms time step. If during the delay period we could decode sample direction from the synaptic efficacies, but not neuronal activity, it would indicate that STP allows for activity-silent maintenance of information.

Sample decoding accuracy using synaptic efficacies (magenta curves, each curve represents decoding from one of twenty networks) was equal to one (perfect decoding) for most of the sample and the entire delay period across all trained networks. In contrast, decoding accuracy using neuronal activity (green curves) decreased to values below 0.4 for all 20 networks by the end of the delay, with 15 networks showing decoding accuracies near chance (1/8) levels (P>0.05, bootstrap, measured during the last 100ms of delay period, see Methods). Thus, for the DMS task, the sample stimulus was perfectly encoded by synaptic efficacies in all 20 networks, and either weakly encoded, or not encoded at all, in neuronal activity.

Although the decoding accuracies provide us with a measure of how much information is stored in neuronal activity and synaptic efficacies, it does not address how the network is using either substrate to solve the task. We wanted to 1) measure how networks used information maintained in neuronal activity and synaptic efficacies to solve the task, and 2) how these contributions relate to the neuronal sample decoding accuracy.

We answered these questions by disrupting network activity or synaptic efficacies during task performance. Specifically, we simulated each trial starting at test onset using the exact same input activity in three different ways: 1) we used the actual neuronal activity and synaptic efficacies taken at test onset as starting points, 2) same as 1, except that we then shuffled the neuronal activity between trials, and 3) same as 1, except that we then shuffled the synaptic efficacies between trials. In each of the three cases, we calculated whether the network output indicated the correct choice.

These results are shown in Figure 2C, comparing neuronal decoding accuracy (x-axis) and task accuracy of the networks (y-axis). Blue, red and cyan circles represent results from the 20 networks using non-shuffled data, shuffled neuronal activity, and shuffled synaptic efficacies, respectively. This allows us to relate decoding accuracy of the sample stimulus (which one could measure in real neurophysiological data sets) with the causal contribution of neuronal and synaptic WM towards solving the task (which is relatively easy to measure in artificial neural network models, but not in neurophysiological experiments).

Neuronal decoding accuracy calculated at the end of the delay was distributed between chance (0.125) and ~0.4, with task accuracy (calculated without shuffling data) consistently > 0.97 (blue circles, right panel, Figure 3A). Networks with strongest delay-period neuronal selectivity suffered the greatest performance loss when their neuronal activities were shuffled (Pearson correlation between neuronal decoding and accuracy after shuffling neuronal activity: R = −0.82, P < 10^−6^, N = 20). Furthermore, networks with the strongest delay-period neuronal selectivity suffered the least performance loss when their synaptic activities were shuffled (correlation between neuronal decoding and accuracy after shuffling synaptic efficacies: R = 0.53, P = 0.015).

**Figure 3.**
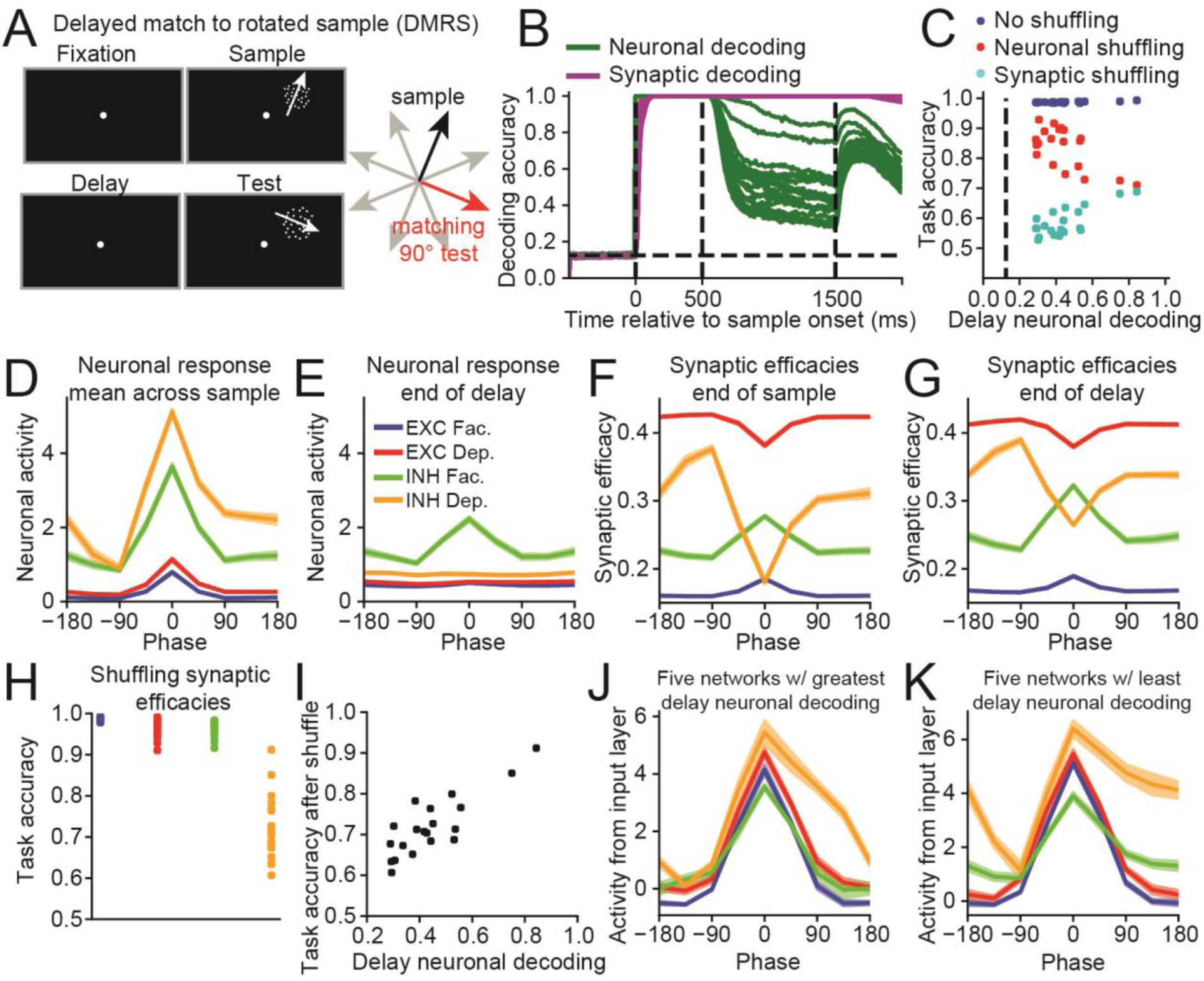
Understanding how networks solve the delayed match-to-rotated sample (DMRS) task. **(A)** The DMRS task is similar to the DMS task (Figure 2A), except that the network was trained to indicate whether the test motion direction was rotated 90° clockwise from the sample motion direction. **(B)** The time course of the sample direction decoding accuracy, calculated using neuronal activity (green curves) or synaptic efficacy (magenta curves) are shown for all twenty networks that successfully solved the DMRS task. The dashed vertical lines, from left to right, indicate the sample onset, offset, and end of the delay period. **(C)** Scatter plot showing the neuronal decoding accuracy measured at the end of the delay (x-axis) versus the behavioral accuracy (y-axis) for all 20 networks trained on this task (blue circles), the behavioral accuracy for the same 20 networks after neuronal activity was shuffled right before test onset (red circles) or synaptic efficacies were shuffled right before test onset (cyan circles). **(D)** The neuronal sample tuning curves were calculated for four groups of neurons (excitatory neurons with facilitating synapses, blue; excitatory neurons with depressing synapses, red; inhibitory neurons with facilitating synapses, green; inhibitory neurons with depressing synapses, orange) from all 20 networks that solved the task. Neuronal activity was averaged across the entire sample period, and the tuning curves were centered around the sample direction that generated the maximum response (i.e. the preferred direction). Error bars (which are small and difficult to see) indicate one SEM. **(E)** Same as (D), except that neuronal activity at the end of the delay period was used to calculate the tuning curves. In order to compare how neural activity evoked during the sample evolves across the delay period, tuning curves were aligned to the preferred directions calculated in (D). **(F)** Same as (D), except that synaptic efficacies at the end of the sample period were used to calculate the tuning curves. As above, the preferred directions are the same as those used in (D). **(G)** Same as (D), except that synaptic efficacies at the end of the delay period were used. **(H)** Task accuracy after shuffling synaptic efficacies at the end of the sample period for each of the four groups of neurons. Each dot represents the accuracy from one network. **(I)** Scatter plot showing the neuronal decoding accuracy measured at the end of the delay period (x-axis) against the task accuracy after shuffling the synaptic efficacies of inhibitory neurons with depressing synapses at the end of the sample period (y-axis). Each dot represents the results of one of the 20 networks. **(J)** Tuning curves showing the activity received from the neurons in the input layer, which was calculated by multiplying the motion direction tuning of the neurons in the input layer with the input-to-recurrent weight matrix. Results were averaged across the 5 networks with the greatest neuronal decoding accuracy during the last 100 ms of the delay period. The four curves thus indicate the mean amount of input each group of recurrent neurons (excitatory or inhibitory, facilitating or depressing) receives from the input layer for each direction. As above, the preferred directions are the same as those used in (D). **(K)** Same as (J), except showing the input tuning curves averaged across the 5 networks with the lowest neuronal decoding accuracy during the last 100 ms of the delay period.

For 14 out of the 15 networks that solved the task using activity-silent WM, in which neuronal decoding accuracy at the end of the delay was not significantly different than chance (P>0.05), shuffling neuronal activity had no significant impact on behavior (P>0.05, bootstrap), and shuffling synaptic efficacies decreased behavioral accuracy to values not significantly greater than chance (P>0.05, bootstrap). Even more drastic perturbations to neuronal activity, such as setting activity for all recurrently connected neurons to zero for the last 100 ms of the delay period, only had a marginal effect on performance (Figure S1), confirming that information maintained in synaptic efficacies, and not neuronal activity, was used to solve the DMS task. Thus, STP allows networks to silently maintain information for the DMS task.

Intuitively, the ability of networks to silently maintain information in WM should depend upon the ratio between the delay duration and the values of STP time constants. We found that networks were unable to solve the DMS task in an activity-silent manner when we decreased the STP time constants to 100 and 500 ms (Figure S2A), or increased the delay to 2500 ms (Figure S2D). In contrast, some networks were still able to solve in activity-silent manner for delay periods of 1500 and 2000 ms (Figure S2B&C). This confirms that our networks are only capable of activity-silent mnemonic encoding when the delay period is not significantly greater than the slow STP time constant.

### Manipulating information

Given that STP can allow networks to silently maintain information in WM, we examined whether it could also allow networks to silently manipulate information. Thus, we repeated the analysis from Figure 2 on 20 networks trained to solve a delayed match-to-rotated (DMRS) sample task, in which the target test motion direction was rotated 90° clockwise from the sample (Figure 3A). Neuronal decoding accuracy for this task (green curves, Figure 3B) was greater than the DMS task (t(38) = 7.22, P < 10^−7^, two-sided, unpaired, t-test, measured during last 100 ms of delay period), suggesting that more information was maintained in neuronal activity compared to networks trained on the DMS task.

Unlike the DMS task, all 20 networks maintained information in a hybrid manner, with significantly elevated synaptic decoding accuracy at the end of the delay compared to chance (P<0.05, bootstrap), and shuffling either neuronal activity or synaptic efficacies significantly decreased behavioral accuracy (P<0.05, bootstrap, Figure 3C). We note that persistent activity exists for this task despite the penalty on high neuronal activity (see Methods).

Consistent with the DMS task, we found that networks with strongest delay-period neuronal selectivity suffered the greatest performance loss when their neuronal activities were shuffled (Pearson correlation coefficient R = −0.65, P = 0.002, N = 20, Figure 3C), and that networks with the strongest delay-period neuronal selectivity suffered the least performance loss when their synaptic activities were shuffled (R = 0.75, P < 0.001).

Although all 20 networks solved the task using persistent activity during the delay, we wondered whether it was still possible that STP was responsible for manipulating sample information. Thus we sought to understand the specific strategy the networks used to solve this task. We examined the mean neuronal responses averaged across the sample for all 20 networks from 4 groups of neurons: excitatory or inhibitory neurons with facilitating or depressing synapses (Figure 3D). We found a striking asymmetry for inhibitory neurons with depressing synapses, in which the sample direction that produced the maximum response was 90° clockwise from the sample direction that produced the weakest response. We can quantify this asymmetry as the difference between the neuronal response 90° clockwise and 90° counterclockwise from the preferred sample direction, and this difference was the greatest for inhibitory neurons with depressing synapses compared to the other three groups of neurons (compared to excitatory neurons with facilitating synapses: t(19) = 8.68, P < 10^−8^; compared to excitatory neurons with depressing synapses: t(19) = 8.26, P < 10^−7^; compared to inhibitory neurons with facilitating synapses: t(19) = 6.69, P < 10^−6^, paired, two-sided, t-tests). This asymmetry in the neuronal activity tuning functions mostly disappeared by the end of the delay period (Figure 3E), but was present in the synaptic efficacies measured at the end of the sample period (Figure 3F), and at the end of the delay period (Figure 3G).

Thus, synaptic efficacies for inhibitory neurons with depressing synapses were at their maximum by the start of the test period on trials in which the sample stimulus was 90° counterclockwise from their preferred direction. If such a sample stimulus is followed by a target test stimulus (90° clockwise from the sample), the total amount of “current” (neuronal response multiplied by synaptic efficacy) these neurons will project upon the rest of the network will be at a maximum. This inhibitory signal can alter the pattern of activity that excitatory neurons project onto the output layer, potentially decreasing the input into the “non-match” output neuron while increasing the input into the “match” output neuron.

Shuffling the synaptic efficacies of inhibitory neurons with depressing synapses at the end of the sample resulted in the lowest task accuracy compared to shuffling efficacies form the other three groups of neurons (compared to excitatory neurons with facilitating synapses: t(19) = − 16.09, P < 10^−11^; compared to excitatory neurons with depressing synapses: t(19) = −14.29, P < 10^−11^; compared to inhibitory neurons with facilitating synapses: t(19) = −12.42, P < 10^−9^, paired, two-sided, t-tests, Figure 3H). Furthermore, networks that maintained the least amount of information in neuronal activity during the delay period were more adversely affected by shuffling their synaptic efficacies (R = 0.85, P < 10^−5^, N = 20, Figure 3I).

We hypothesized that the asymmetric tuning of inhibitory neurons with depressing synapses was the result of the connection weights from the input layer. Thus, we calculated the tuning curves for the current (neuronal activity times the connection weight) neurons receive from the input layer. For the five networks that maintained the greatest amount of information in neuronal activity during the delay (Figure 3J), their tuning curves were again asymmetric, but the direction producing the weakest input was not clearly 90° counterclockwise from the direction producing the greatest input. In contrast, for the 5 networks that maintained the least amount of information in neuronal activity during the delay (Figure 3K), their tuning curves were strongly asymmetric with the direction producing the weakest input 90° counterclockwise from the direction producing the greatest input.

We confirmed that these results were not specific to this one task by repeating our analysis on 20 networks with a DMRS task using a 90° *counterclockwise* rule (Figure S3). These results were the mirror image of the results shown in Figure 3. We also repeated our analysis on a delayed match-to-category (DMC) task, in which the networks had to indicate whether sample and test stimuli belonged to the same learned categories^32^. We found that the networks performed the manipulation (i.e. categorized the stimulus) by adjusting connection weights from the input layer (Figure S4). Given that we imposed an incentive for the networks to reduce neuronal activity, our results suggest that networks will attempt to perform the required manipulation of the sample stimulus by learning specific connection weights from the input layer if possible.

In summary, STP was not directly involved in manipulating the sample stimulus in the DMRS task. Rather, networks performed the manipulation by adjusting connection weights from the input layer. However, this analysis reveals that STP can generate a prospective code^33^, facilitating the proper network response to upcoming test stimuli.

### Manipulating information during the WM delay period

We considered whether STP can play a more active role manipulating information if the manipulation cannot be performed by adjusting input connection weights. This could be accomplished by forcing the manipulation to occur after the sample stimulus is extinguished— when the input layer no longer remains active. We trained networks to solve a delayed cue task (Figure 4A), in which a cue was presented between 500 and 750 ms into the delay, instructing the network whether identical sample and test directions were a match (DMS), or whether a test direction rotated 90^0^ clockwise from a sample was the target (DMRS).

**Figure 4.**
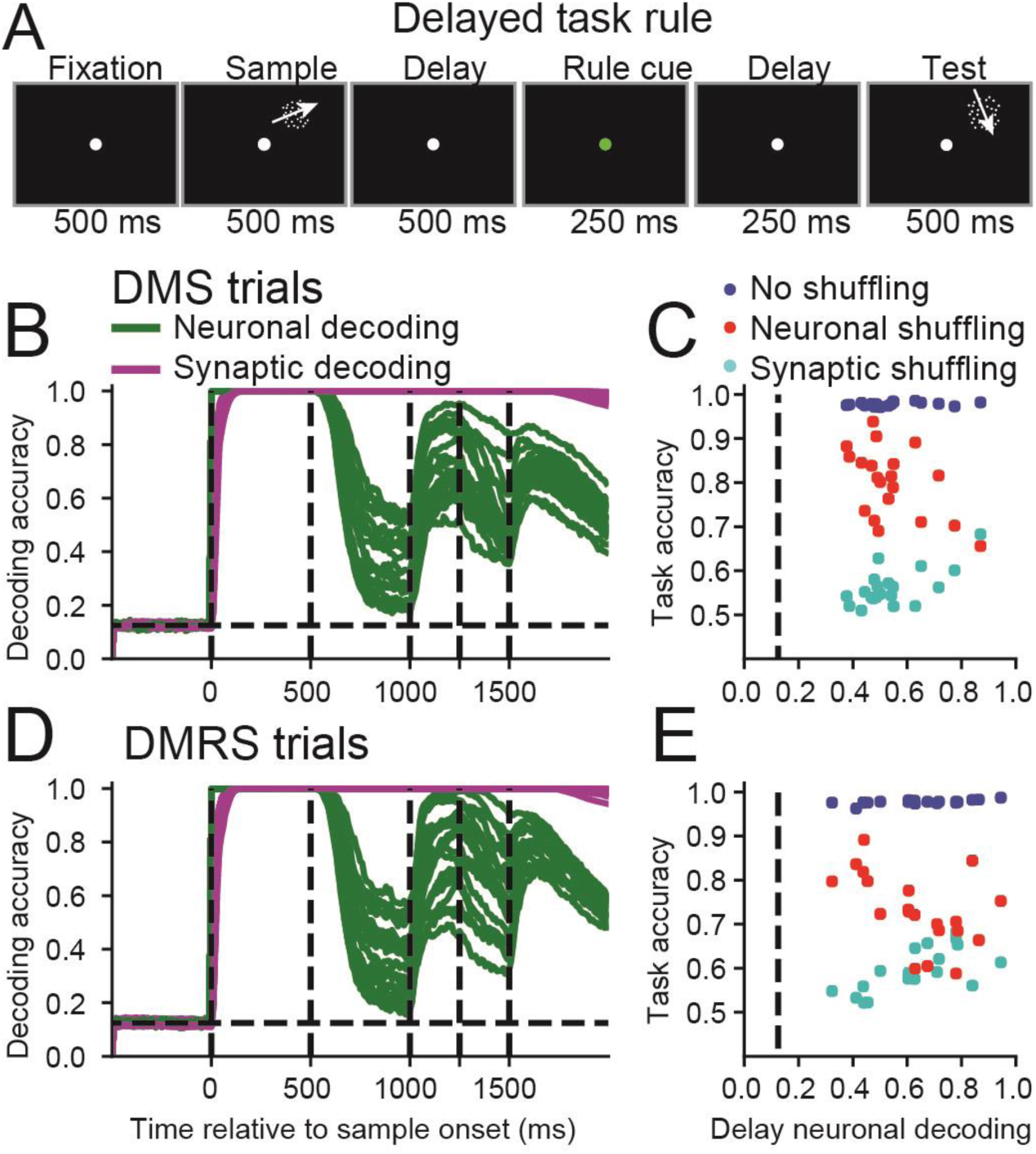
Neuronal and synaptic mnemonic encoding for tasks that involve manipulating the contents of working memory. **(A)** This task was similar to the DMS and DMRS tasks, except that a rule cue from 500 to 750 ms into the delay period indicated to the network whether to perform the DMS or the DMRS task. **(B)** Similar to Figure 2B, except showing the decoding results for DMS trials for networks trained to solve the delayed rule task. The dashed vertical lines, from left to right, indicate the sample onset and offset, the rule cue onset and offset, and end of the delay period. **(C)** Similar to Figure 2C, except showing the shuffle analysis for DMS trials for networks trained to solve the delayed rule task. **(D-E)** Same as (B-C), except showing the decoding results and shuffle analysis for DMRS trials (in which matching sample and test stimuli are rotated by 90°).

We found that none of the 20 networks silently-maintained information for either trial type as the sample neuronal decoding accuracy never decreased to chance levels (P > 0.05, bootstrap) during the delay for either DMS (green curves, Figure 4B) or DMRS trials (Figure 4D). Thus, these networks manipulate information in WM (at least) partly using persistent neuronal activity.

We next examined the contribution of neuronal activity and synaptic efficacies toward solving the task by shuffling neuronal activity or synaptic efficacies immediately prior to test onset. Consistent with Figures 2&3, task accuracy after shuffling activity (red circles) was negatively correlated with neuronal decoding accuracy during the end of the delay (DMS: R = −0.51, P = 0.021, N = 20, Figure 4C; DMRS: R = −0.49, P = 0.028, Pearson correlation coefficient, Figure 4E), implying that for networks showing weak neuronal decoding, shuffling neuronal activity had little impact on behavior. Furthermore, behavioral accuracy after shuffling synaptic efficacies (cyan circles) was positively correlated with neuronal decoding accuracy during the end of the delay (DMS: R = 0.66, P = 0.002; DMRS: R = 0.71, P < 0.001), implying that networks showing weak neuronal decoding were strongly affected by shuffling synaptic efficacies. Furthermore, shuffling neuronal activity or synaptic efficacies significantly decreased behavioral accuracy (P < 0.05, bootstrap) in all 20 networks for both tasks.

Thus, networks required neuronal activity to manipulate information maintained in WM.

However, consistent with previous tasks, information was maintained in a hybrid state during the delay.

### Controlling the representation of information

Although STP did not silently manipulate information during the tasks considered so far, we wondered if it could allow for subtler manipulations of information in a silent manner. For example, neural circuits *in vivo* are occasionally required to represent behaviorally-relevant information in a different format compared to irrelevant information^34^. Thus, we trained networks on a task that required networks to control how information is represented in the network: the A-B-B-A task^35^ (Figure 5A). Here, networks were presented with a sample stimulus followed by three sequential test stimuli, and had to indicate whether each test stimulus matched the sample. Importantly, if a test stimulus did not match the sample, there was a 50% probability that the test stimulus would be repeated on the subsequent test presentation. This forces the network to encode sample and test stimuli in different ways: if the sample and test were represented in similar manners, then the network could not distinguish between a test stimulus that matched the sample (match event), compared to a test stimulus that matched the previous test (non-match event). We provide evidence that the networks represent sample and test stimuli in different formats below.

**Figure 5.**
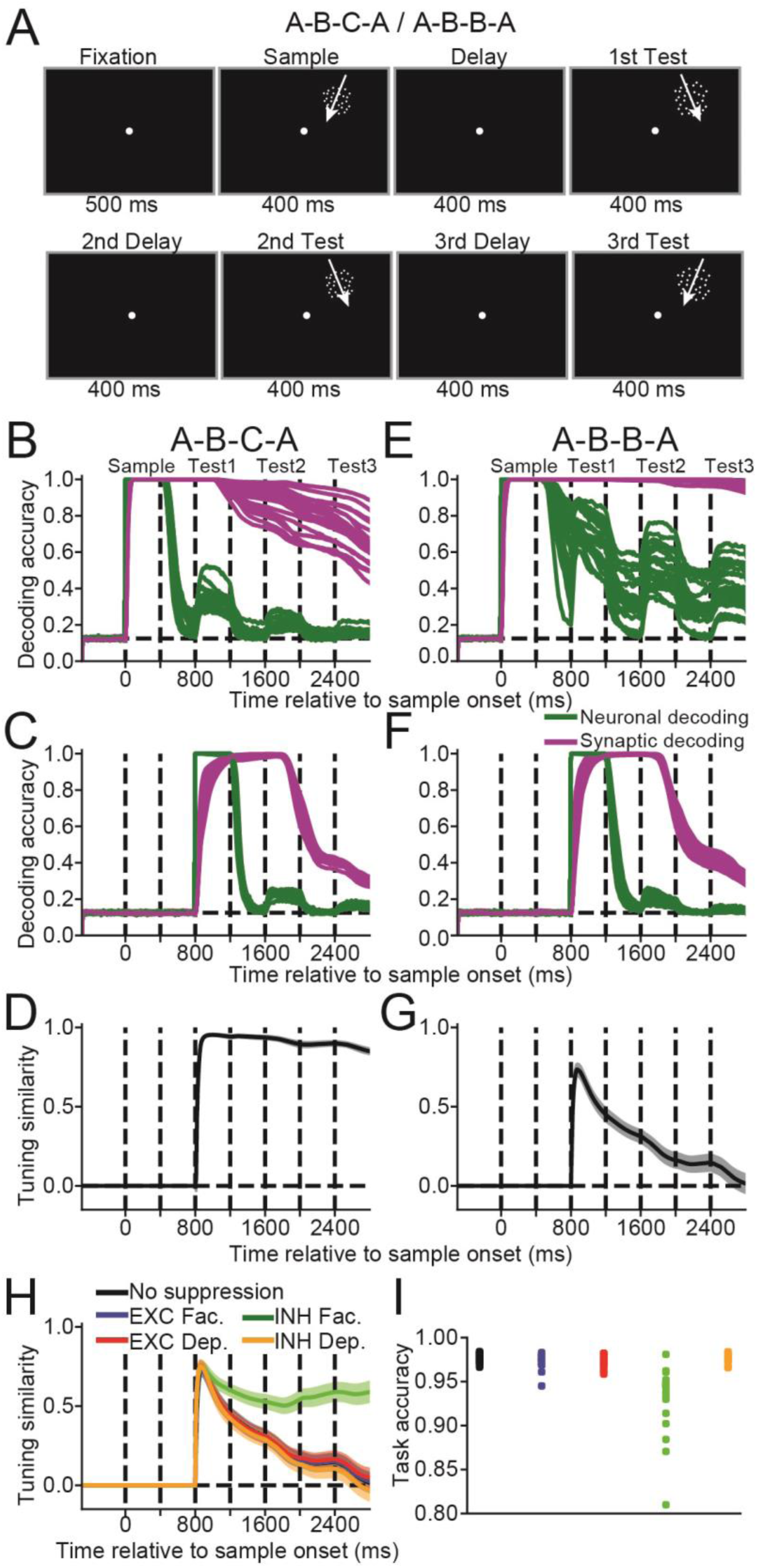
Neuronal and synaptic mnemonic encoding for tasks that require controlling how information is represented. **(A)** In the A-B-B-A and A-B-C-A task, the network was presented with a 400 ms motion direction stimulus, followed by three 400 ms motion direction test stimuli, in which all stimuli were separated by a 400 ms delay. The network was trained to indicate whether each test stimulus matched the motion direction of the sample stimulus. For test stimuli two and three in the A-B-B-A task, there was a 50% chance that a non-matching test stimulus would have the motion direction as the previous non-matching test stimulus. For the A-B-C-A task, non-matching test stimuli never matched any previous test stimuli. **(B)** Similar to Figure 2B, the time course of the sample direction decoding accuracy, calculated using neuronal activity (green curve) or synaptic efficacy (magenta curve) are shown for all 20 networks trained on the A-B-C-A task. The dashed lines indicate, from left to right, the sample onset, the sample offset, and the test onset and offset for the three sequential test stimuli. **(C)** Similar to (B), except showing the decoding accuracy of the first test stimulus. **(D)** The time course of the tuning similarity index (the weighted dot-product between the preferred sample motion direction and the preferred first test motion direction averaged across all synaptic efficacies), averaged across all 20 networks. **(E-G)** Similar to (B-D), except for the A-B-B-A task. **(H)** The mean tuning similarity index is shown for the A-B-B-A task, after suppressing neuronal activity for 200 ms before the first test onset, from four different groups of neurons (excitatory neurons with facilitating synapses, blue curve; excitatory neurons with depressing synapses, red curve; inhibitory neurons with facilitating synapses, green curve; inhibitory neurons with depressing synapses, cyan curve), and for the case when activity was not suppressed (black curve). **(I)** Behavioral task accuracy for the cases when no activity was suppressed (black dots), and after suppressing activity from the four groups of neurons described in (H). Each dot represents the task accuracy from one network.

As a control, we also trained networks on an A-B-C-A version of the task, which was similar to the A-B-B-A task, except that non-matching test stimuli were never repeated during a single trial, so that the network is not forced to represent sample and test stimuli in a different manner.

For the A-B-C-A task, most networks appeared to solve the task using information in synaptic efficacies, as few networks encoded sample information in neural activity throughout the entire trial. We found that sample decoding using neuronal activity decreased to chance levels (P>0.05, bootstrap) for 1 of 20 networks during the last 100 ms of the first delay, 15 of 20 networks during the second delay, and 18 of 20 networks for the third delay (green curves, Figure 5B). In contrast, sample decoding using synaptic efficacies (magenta curves) remained significantly above chance (P<0.05) throughout the entire trial for all 20 networks (values ranging from ~0.55 to 1.0). We note that decoding accuracy appeared relatively lower for this task because of how we performed the calculation (see Supplementary Information).

Since networks were trained to compare each test direction to that of the previous sample, it made sense that sample information was represented throughout the trial. However, do networks also maintain test information in WM, which is only behaviorally relevant during test presentation? We found that decoding accuracy for the first test stimulus using neuronal activity (green curves) was perfect (1.0) for all networks during test presentation, before rapidly dropping to chance levels (P>0.05) for all 20 networks by the third delay period (Figure 5C).

Test decoding using synaptic efficacies (magenta curves) was near perfect (~1.0) for all networks during the later stage of the first test presentation, the subsequent delay, and into the second test presentation. Thus, both the sample and first test motion directions were mnemonically encoded by the network during presentation of the second test stimulus. This could be problematic if the network had to distinguish between cases where the second test stimulus matched the sample vs the first test. However, this was not a problem for the A-B-C-A task, as non-matching test stimuli were never repeated.

As discussed above, the network was under no pressure to represent sample and test stimuli differently during the A-B-C-A task, which we confirmed using a tuning similarity index^10^, which was the weighted dot-product between the preferred sample first test directions averaged across all synaptic efficacies (Figure 5D see Methods). A value of +1 indicates that synaptic efficacies are identically tuned to sample and test stimuli, 0 indicates no correlation between the two, and −1 indicates that synaptic efficacies prefer opposite sample and test directions. As expected, the tuning similarity index was >0.9 for the first and second test periods. This implies that for each synapse, the preferred sample tended to be similar to the preferred test— suggesting similar representation of sample and test information by synaptic efficacies.

We then repeated these analyses for the A-B-B-A task, in which subsequent non-matching test stimuli were repeated 50% of the time. As with the previous task, we decoded the sample motion direction using neuronal activity (green curves, Figure 5E) and synaptic efficacies (magenta curves). In contrast to the A-B-C-A task, sample decoding accuracy using neuronal activity in the A-B-B-A task remained above chance (P<0.05, bootstrap) during all three delay periods for 19 out of 20 networks. Furthermore, sample decoding accuracy using synaptic efficacies remained close to 1.0 throughout the trial.

Consistent with the A-B-C-A task, decoding accuracy for the first test stimulus using neuronal activity (green curves, Figure 5F) was perfect (1.0) during the first test presentation before rapidly falling to chance (P>0.05) levels after test stimulus offset. Also consistent with the A-B-C-A task, decoding accuracy for the first test stimulus using synaptic efficacies (magenta curves) was also near perfect (~1.0) for all networks during the later stages of the first test presentation, the subsequent delay, and into the second test presentation. As alluded to above, it is potentially problematic that the sample and the first test stimulus were both encoded by the network during the second test presentation, as the network must distinguish between cases where the second test matched the sample (match event), or the first test (non-match event).

We hypothesized that the sample and the first test stimuli must be represented in different formats for networks to accurately solve the task. To confirm this, we calculated the tuning similarity index for all 20 networks. In contrast to the A-B-C-A task (Figure 5D), in which the mean tuning similarity index was 0.92 ± 0.05 (SD) during the second test period, the similarity index for the A-B-B-A task decreased to 0.23 ± 0.18 (t(38) = 16.21, P < 10^−17^, unpaired two-sided, t-test, Figure 5G). Thus, for synapses in the A-B-B-A task, preferred sample stimuli tended to differ from preferred test stimuli, implying different encoding formats. By so doing, the network can theoretically distinguish between cases in which subsequent test stimuli match the sample (match event) vs earlier test stimuli (non-match event)

We next asked how the network was able to represent the sample and first test stimuli in different formats. We hypothesized that delay-period persistent neuronal activity was necessary to encode the first test stimulus in a different format than the sample. Thus, we suppressed neuronal activity from four different groups of neurons for the 200 ms period prior to the first test, and re-calculated the tuning similarity index (Figure 5H). Suppressing activity from inhibitory neurons with facilitating synapses increased the mean tuning similarity index, measured during the second test period (no suppression = 0.232 ± 0.18, suppression = 0.516 ± 0.19, green curve, t(19) = 5.83, P < 10^−4^, paired, two-sided, t-test). Furthermore, suppressing the activity of inhibitory neurons with facilitating synapses decreased behavioral accuracy significantly more than suppressing the activity of any of the other three groups of neurons (compared to excitatory neurons with facilitating synapses: t(19) = −5.71, P < 10^−4^; compared to excitatory neurons with depressing synapses: t(19) = −5.39, P < 10^−4^; compared to inhibitory neurons with depressing synapses: t(19) = −5.33, P < 10^−4^, paired, two-sided, t-tests, Figure 5I). Thus, neuronal activity from these neurons is required to manipulate information in short-term memory, increasing task performance.

In summary, delay-period neuronal activity from inhibitory neurons with facilitating synapses was required to represent sample and test information in different formats, and suppressing this activity resulted in similar sample and test representations along with a decrease in behavioral accuracy. Interestingly, this contrasts with the results of the DMRS task (Figure 3), in which inhibitory neurons with depressing synapses played a critical role in representing task-relevant information. Thus, while sample information maintained in synaptic efficacies was primarily used by the networks to solve the A-B-C-A and A-B-B-A tasks, neuronal activity was required to control how information was represented.

### Attending to specific memoranda

Recent studies suggest that silently-maintained information can be reactivated either by focusing attention towards the memorandum^19^ or by probing the neural circuits involved^20^. In a final experiment, we examined how STP supports short-term maintenance of information that is either attended or unattended. We trained networks on a dual-sample delayed matching task (Figure 6A) that roughly follows the study by Rose et al.^19^. The network was trained to maintain two sample directions (presented simultaneously in two different locations) in WM, followed by two successive cues and test stimuli, in which the cue indicated which of the two samples was relevant for the upcoming test. In this setup, it is possible that stimuli that were not cued as relevant for the first test stimulus, could still be cued as relevant for the second test stimuli. Thus, stimulus could switch from being unattended to attended.

Across the 20 networks, the mean sample decoding accuracy using neuronal activity was significantly greater when the sample was attended (blue curve) compared to unattended (red curve), during the last 100 ms of the first and second delay periods (first delay: Figure 6B, left panel, attended = 0.46 ± 0.25, unattended = 0.35 ± 0.19, t(39) = 4.02, P < 0.001; second delay: right panel, attended = 0.34 ± 0.21, unattended = 0.22 ± 0.10, t(39) = 5.49, P < 10^−5^, paired, two-sided, t-tests). Sample decoding accuracy using synaptic efficacies was near perfect (~1.0) for both the attended (black curves) and unattended (yellow curves) conditions. Thus, the attended memoranda were more strongly represented in neural activity compared to the unattended memoranda.

The study by Rose et al. found that unattended information that was silently maintained could be reloaded back into neuronal activity after it was attended^19^. Similarly, we found that neural decoding significantly for stimuli that were unattended after the first cue became the focus of attention after the second cue (blue dots, Figure 6C), compared to stimuli that remained unattended to (red dots, t(39) = 5.30, P < 10^−5^, paired, two-sided, t-test)).

Although sample decoding accuracy using neuronal activity was not significantly greater than chance for many networks, the rule cue indicating which stimulus to attend to was encoded and maintained in neuronal activity (neuronal decoding accuracy for rule cue #1 is indicated by the dashed green curve, and rule cue #2 is indicated by the solid green curve, Figure 6D). Thus, while sample information can be silently maintained, allocating attention to either memoranda requires neuronal activity.

**Figure 6.**
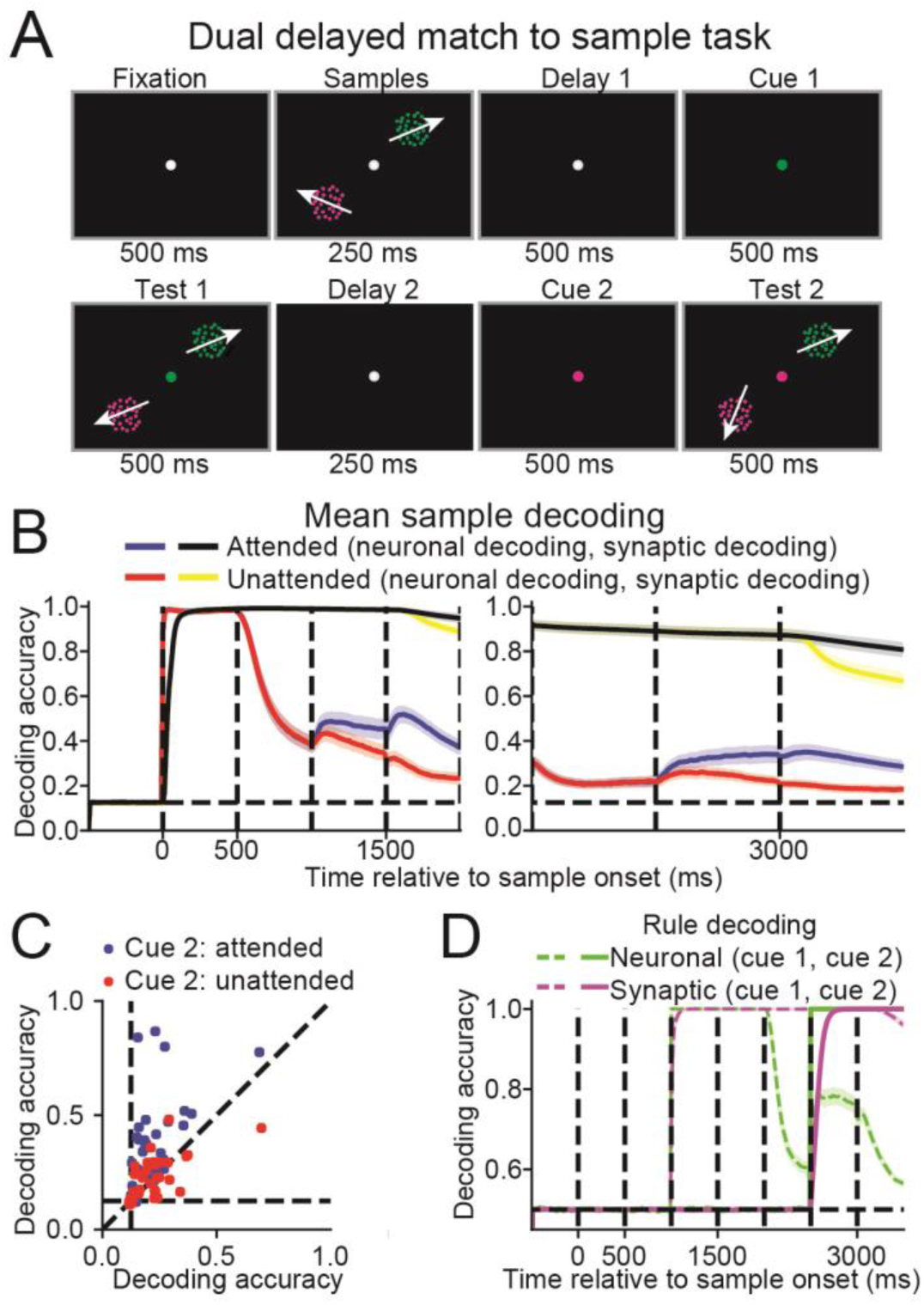
Neuronal and synaptic mnemonic encoding of multiple stimuli. **(A)** In the dual DMS task, two sample stimuli were simultaneously presented for 500 ms. This was followed by a 1000 ms delay period in which a cue appeared halfway through, and then two simultaneous test stimuli for 500 ms. The cue indicated which of the two sample/test pairs were task-relevant. Another 1000 ms delay and 500 ms test period was then repeated, in which a second cue again indicated which of the two sample/test pairs was task-relevant. **(B)** Neuronal decoding accuracy for the attended (blue curve) and unattended (red) stimuli, and the synaptic decoding accuracy for the attended (black) and unattended (yellow) stimuli, are shown from trial start through the first test period (left panel), and the second delay and test periods (right panel). **(C)** Scatter plot showing the neuronal decoding accuracy, measured from 100 to 0 ms before the second cue (x-axis) against the neuronal decoding accuracy, measured from 400 to 500 ms after second cue (y-axis). Blue dots represent stimuli that were unattended after the first cue, and attended after the second cue, and red dots represent stimuli that were not attended to after the first and second cues. **(D)** The neuronal (green) and synaptic (magenta) rule decoding accuracy. The dashed lines indicate the decoding accuracy of the first cue, and the solid lines indicate the decoding accuracy of the second cue.

### Manipulating information and persistent neuronal activity

While STP can allow networks to silently maintain information, persistent neuronal activity during the delay period was observed in all the tasks that involved manipulating information (DMRS, delayed cue, ABBA, attention in the dual-sample task). We wondered if the level of manipulation required by the task was correlated with the level of persistent activity. This could be of special interest as varying levels of persistent activity have been observed between different tasks^10–16^.

We reasoned that for tasks that did not require manipulation, the network should represent the sample stimulus in fundamentally the same way during all trial epochs (e.g. early sample vs late delay). One caveat is that networks trained on tasks that did not require manipulation (e.g. DMS) did not represent the sample in neuronal activity at the end of the delay. However, in all cases the sample stimulus was represented in synaptic efficacies, with high decoding accuracy throughout the delay. Thus, we quantified the level of manipulation for each task by computing the similarity (same as used in Figure 5, see Methods) between the neuronal response during the first 100 ms of sample onset, and the synaptic efficacies during the last 100 ms of the delay. We subtracted this value from 1.0, so that a manipulation near 0 implies that early-sample tuning was similar to late-delay synaptic tuning. To boost statistical power, we included two additional delayed match-to-rotated sample tasks in which the target test direction was 45° clockwise from the sample direction (DMRS45) and 180° from the sample direction (DMRS180).

We found that the level of manipulation was correlated with the level of stimulus-selective persistent activity (i.e. neuronal decoding accuracy) measured during the end of the delay (Spearman correlation coefficient R = 0.79, P = 0.006, N = 10, Figure 7). This suggests that tasks requiring greater manipulation require greater levels of persistent activity. Thus, different levels of manipulation could partly explain the observation that persistent neuronal activity varies markedly between tasks.

**Figure 7.**
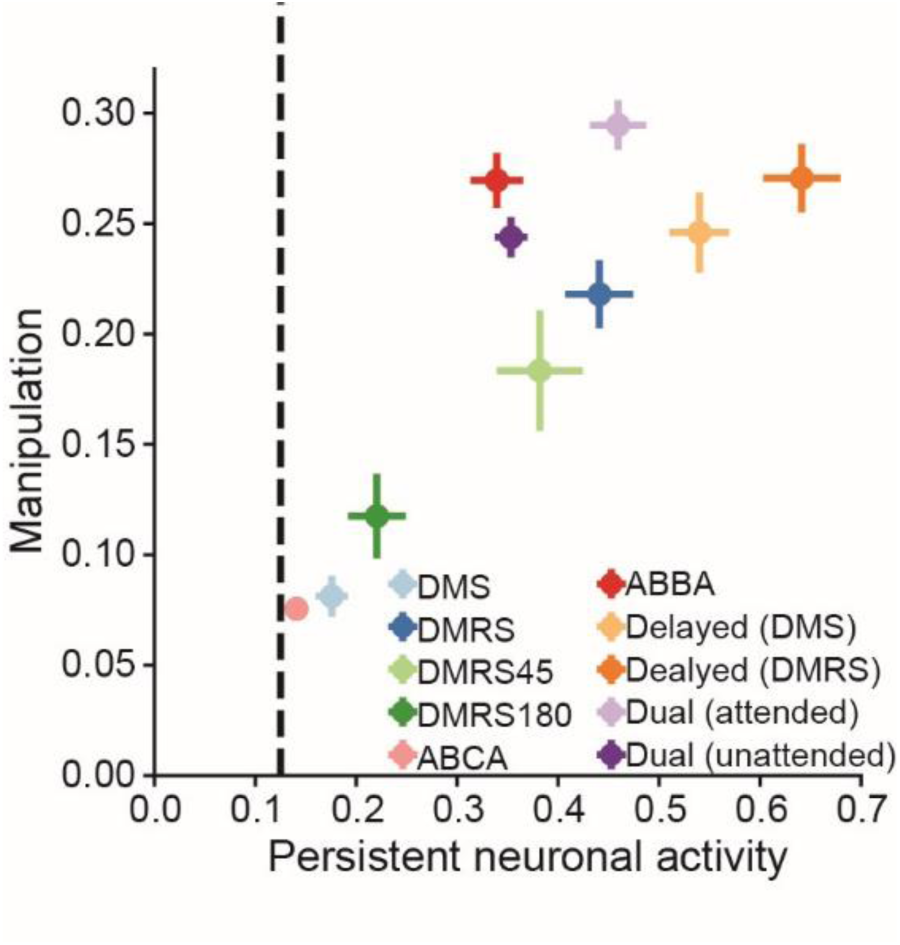
The relationship between the manipulation required by the task and the level of stimulus-selective persistent neuronal activity. Scatter plot shows the level of stimulus-selective persistent neuronal activity, measured as the neuronal decoding accuracy during the last 100 ms of the delay period (x-axis), versus the level of manipulation, which was equal to 1 minus the tuning similarity between the neuronal tuning during the first 100 ms of the sample period, and the synaptic tuning during the last 100 ms of the delay period (y-axis). Each dot represents the average values across 20 networks trained on one specific task, or the across one trial type from one specific task (i.e. the DMS and DMRS trials from the delayed rule task, and the attended and unattended trials from the dual-sample delayed matching task). Errors bars indicate one SEM. Dashed vertical line indicates chance level decoding.

## Discussion

We examined whether STP can support the activity-silent manipulation of information, and whether it could help explain previous observations that different tasks evoke different levels of persistent activity. We found that while STP can silently support the short-term maintenance of information, it cannot support manipulation of information without persistent neuronal activity. Furthermore, we found that tasks that required more manipulation also required more persistent activity, giving insight into why the strength of persistent neuronal activity varies markedly between different tasks.

### Variation in persistent neuronal activity in vivo

Over the last several decades, monkey electrophysiology experiments^2–6^, and later human imaging studies^36^, have supported the idea that information in WM is maintained in stimulus-selective persistent neuronal activity during memory-delay periods of behavioral tasks. However, this viewpoint has evolved, as various studies now suggest that persistent neuronal activity might not always reflect information maintenance, but can reflect control processes required to manipulate remembered information into appropriate behavioral responses^14^.

It is often unclear whether persistent neural activity reflects the maintenance or the manipulation of the stimulus. For example, neural activity in the frontal and parietal cortices mnemonically encode stimulus location in a memory delayed saccade task^2,4^. However, recent studies that have dissociated the stimulus location from the upcoming saccade location have shown that activity in frontal cortex initially encodes the location of the recent stimulus (retrospective code), before its representation shifts towards encoding the planned saccade target (prospective code) later in the delay^37^.

In another study, Mendoza Halliday *et al*.^38^ showed robust persistent activity in the medial superior temporal (MST) area during a motion DMS task. This initially appears at odds with the results of our current study, and our past work showing little or no persistent activity in the lateral intraparietal area (LIP), an area downstream of MST, also during a motion DMS task^10,16^. However, the Mendoza Halliday et al. task presented sample and test stimuli in different retinotopic locations. This forces MST to represent the sample and test stimuli using two different pools of neurons, eliminating the possibility that synaptic efficacy changes through STP driven by the sample could be directly compared to the test stimulus activity. This also forces the monkey to translate information from the sample location to the location of the test stimulus. Moreover, while we only observed weak delay-period direction encoding in area LIP during the DMS task^10,16^, we found that after the monkeys underwent extensive categorization training using the same stimuli, delay-period categorical encoding become highly robust^16^.

In another example, prefrontal cortex (PFC) was shown to mnemonically encode color in a change-detection task when six, highly distinguishable, colors are used^39^, but color-selective persistent activity was not evident in PFC when the subject had to detect a change amongst a continuum of 20 colors^12^. This might suggest that PFC activity can encode a categorical representation of the stimulus, but not a precise representation of stimulus features.

These results suggest that persistent neuronal activity emerges as a result of manipulating the stimulus representation into another format such as a categorical code or a planned behavioral response, or through the allocation of attention. In contrast, studies have found that persistent neuronal activity is weak^40^ to non-existent^38,41^ in sensory areas, which are expected to be more involved in stimulus representation rather than manipulation. Despite the relative lack of persistent neuronal activity, studies have found that stimulus-selective information can still be decoded from local-field potentials^38^ and from blood-oxygenation levels^42^, suggesting that subthreshold top-down signals from frontal areas may coordinate with posterior areas to maintain information in short-term memory.

In summary, these studies suggest that tasks that require greater manipulation of the memoranda evoke greater levels of persistent neuronal activity across a greater number of cortical areas. Although other factors undoubtedly influence the level of persistent neuronal activity, such as circuit properties^10,43^ and other task-related variables^44^, we observed a similar correlation between the level of manipulation and the level of persistent neuronal activity in our network models (Figure 7). Thus, our results suggest that the variability in the level of persistent neuronal activity observed between different tasks is partially the result of varying levels of stimulus manipulation required between tasks.

### Comparison to other artificial neural network architectures

Until recently, it was difficult to train RNNs to solve tasks involving long-term dependencies, in which the desired network outputs depend on inputs separated by long temporal delays. In the last several years, long short-term memory (LSTM)^45^ and more recently, gated recurrent unit (GRU)^46^ architectures have shot to prominence by allowing RNNs to solve a variety of tasks such as language modelling and translation that involve very long temporal delays. These architectures work by allowing networks to determine which information to maintain across time, and which information to update.

We noticed that RNNs without STP either failed to accurately solve the task, or required many more training epochs. This was true even when setting the penalty on neuronal activity to zero (Figure S5). This difficulty in solving the tasks was partly because neurons in our networks never connected onto themselves, which could otherwise help neuron maintain information in short-term memory. That said, STP, with its relatively long time constants, greatly facilitated training on our set of WM based tasks. This highlights how adding network substrates with long time constants, without necessarily making these time constants flexible, can significantly decrease training time on tasks with long-term time dependencies. More generally, it also highlights how incorporating neurobiologically-inspired features to artificial neural networks is a promising strategy for advancing their capabilities.

A recent study proposed a different recurrent network architecture in which activity was temporarily maintained in “fast weights”, which operated on a slower timescale than changes in neural activity, but faster than changes in synaptic weights^47^. Adding this different time constant allowed the network to more accurately solve tasks that required information maintenance across input sequences. These fast weights are reminiscent of the dynamic synapses regulated by STP considered in this study, and further suggest how adding a range of effective time constants to RNN models can improve performance on tasks that require maintaining information across different time scales.

### Understanding strategies employed by artificial neural networks

A key differentiating feature of RNNs compared to biological networks is that all connection weights and activity is known, facilitating any attempt to understand how the network solves various tasks. This has allowed researchers to describe how delayed association in RNNs can be performed by transient dynamics^25^, how simple dynamical systems consisting of line attractors and a selection vector can underlie flexible decision-making^23^, how RNNs can encode temporally-invariant signals^26,27^, and how functional clustering develops when RNNs learn multiple tasks^48^.

While modelling studies cannot directly replace experimental work, they can be highly advantageous when obtaining the necessary experimental data is not feasible. Such was the case with our study, in which current technology does not allow us to directly measure the information maintained in synaptic efficacies or their contribution towards the behavioral decision. Thus, these modelling studies can serve as an excellent complement to experimental work, allowing researchers to rapidly generate novel hypothesis regarding cortical function that can later be tested when technology better allows for experimental verification. Lastly, the discovery of novel mechanisms found *in silico* can be fed back into the design of network models, potentially accelerating the development of machine learning algorithms and architectures. We believe that this synergy between experimental neuroscience and neural network modelling will only strengthen in the future.

## Acknowledgements

This work was supported by National Institutes of Health R01EY019041 and R01MH092927, and National Science Foundation Career Award NCS 1631571.

## Methods

### Code and data availability

Data sharing is not applicable to this study since no experimental datasets were generated or analyzed, but the code used to train, simulate and analyze network activity is available at https://github.com/nmasse/Short-term-plasticity-RNN.

### Network models

Neural networks were trained and simulated using the Python machine learning framework TensorFlow^49^. Parameters used to define the network architecture and training are given in Table 1. All networks consisted of 36 or 48 input neurons (whose firing rates are represented as u(*t*)) that projected onto 100 recurrently connected neurons (whose firing rates are represented as r(*t*)), which in turn projected onto 3 output neurons (whose firing rates are represented as z(*t*)) (Figure 2A).

The activity of the recurrent neurons was modelled to follow the dynamical system^48^:
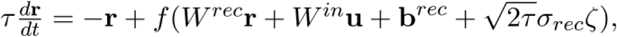

where τ is the neuron’s time constant, *f*(.) is the activation function, *W^rec^* and *W^in^* are the synaptic weights between recurrent neurons, and between input and recurrent neurons, respectively, b is a bias term, ζ is independent Gaussian white noise with zero mean and unit variance applied to all recurrent neurons, and *σ*_r_*_ec_* is the strength of the noise. To ensure that neuron’s firing rates were non-negative and non-saturating, we chose the rectified linear function as our activation function: *f*(*x*) *= max*(*0,x*).

The recurrent neurons project linearly to the output neurons. The activity of the output neurons, *z*, was then normalized by a softmax function such that their total activity at any given time point was one:

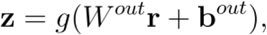

where *W^out^* are the synaptic weights between the recurrent and output neurons, and *g* is the softmax function:

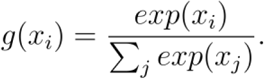

To simulate the network, we used a first-order Euler approximation with time step Δ*t*:

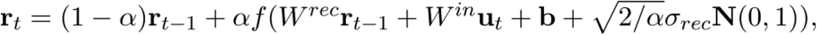

where α *=* Δ*t/r* and N(0, 1) indicates the standard normal distribution.

To maintain separate populations of excitatory and inhibitory neurons, we decomposed the recurrent weight matrix, *W^rec^* as the product between a matrix for which all entries are non-negative, *W^rec:+^* whose values were trained, and a fixed diagonal matrix, *D*, composed of 1s and −1s, corresponding to excitatory and inhibitory neurons, respectively^30^:

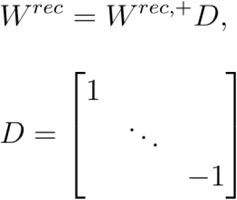

Initial connection weight values were randomly sampled from a Gamma distribution with shape parameter of 0.25 and scale parameter of 1^30^. We note that training networks to accurately solve the tasks appeared somewhat faster when initializing connection weights from a gamma distribution compared to a uniform distribution (data not shown). Initial bias values were set to 0.

Networks consisted of 36 motion direction tuned input neurons, and for the rule switching tasks (i.e. DMS+DMRS and dualDMS tasks), an additional 12 rule tuned neurons. The tuning of the motion direction selective neurons followed a Von Mises’ distribution, such that the activity of the input neuron *i* at time *t* was

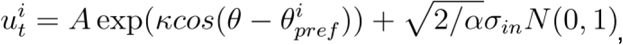

where *θ* is the direction of the stimulus, 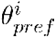 is the preferred direction of input neuron *i, k* was set to 2, and *A* was set to 4*/exp*(*k*) when a stimulus present (i.e. during the sample and test periods), and was set to zero when no stimulus was present (i.e. during the fixation and delay periods). For the dualDMS task, 18 motion direction tuned input neurons represented the first stimulus position, and the second represented the second stimulus position.

The 12 rule tuned neurons for the DMS+DMRS and dualDMS tasks had binary tuning, in which their activity was set to 2 (plus the Gaussian noise term) for their preferred rule cue, and 0 (plus the Gaussian noise term) for all other times. The number of rule tuned neurons was arbitrarily chosen, and has little impact on network training.

### Network training

RNNs were trained based on techniques previously described^30,48^. Specifically, the goal of training was to minimize 1) the cross-entropy between the output activity and the target output, and 2) the mean L2-norm of the recurrent neurons’ spike rate. The target output was a length 3 one-hot vector, in which the first element was equal to one for all times except the test period, the second element was one when the test stimulus matched the sample, and the third element was one when the test stimulus did not match the sample. Specifically, the loss function at time *t* during trial *i* is

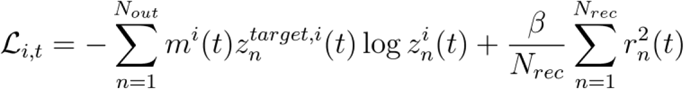

where *β* controls how much to penalize neuronal activity of the recurrent neurons, and *m^i^*(*t*) is a mask function, with values of zero or one, that determine which portions of the trial to include in the loss function. To give the network adequate time to form a match or non-match decision, we set the mask function to zero for the first 50 ms of the test period, and one for all other times. The total loss function is then

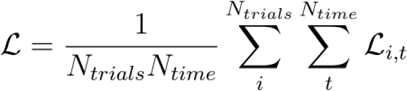

During training, we adjusted all parameters (*W^in^*, *W^rec,+^, W^out^, b^rec^*, *b^out^*) using the Adam^50^ version of stochastic gradient descent. The decay rate of the 1st and 2nd moments were set to their default values (0.9 and 0.999, respectively).

### Short-term synaptic plasticity

The synaptic efficacy between all recurrently connected neurons was dynamically modulated through short-term synaptic plasticity (STP). For half of the recurrent neurons (40 excitatory and 10 inhibitory), all projecting synapses were facilitating, and for the other half of recurrent neurons, all projecting synapses were depressing. Following the conventions of Mongillo et al.^18^, we modelled STP as the interaction between two values: *x*, representing the fraction of neurotransmitter available, and *u*, representing the calcium concentration in the presynaptic terminal. Presynaptic activity acts to increase the calcium concentration inside the presynaptic terminal, increasing synaptic efficacy. However, presynaptic activity decreases the fraction of neurotransmitter available, leading to decreasing efficacy. These two values evolve according to:

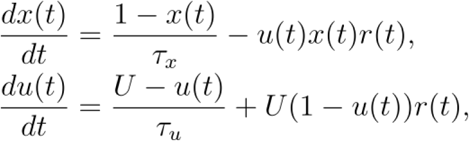

where *r*(*t*) is the presynaptic activity at time *t*, *τ_x_* the neurotransmitter recovery time constant, and *τ_u_* is the calcium concentration time constant. The amount of input the postsynaptic neuron receives through this one synapses at time *t* is then

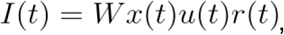

where *W* is the synaptic efficacy before STP is applied. For depressing synapses, the neurotransmitter recovery time constant was much longer compared to the calcium concentration time constant, whereas the opposite was true for facilitating synapses.

### Population decoding

Similar to our previous studies^10,16^, we quantified the strength of stimulus encoding by measuring how accurately we could decode the motion direction using multiclass support vector machines (SVMs). In this approach, we trained linear, multiclass SVMs to classify the motion direction using the neuronal activity of the 100 recurrent neurons, or the synaptic efficacies from the same 100 recurrent neurons, at each time point. The synaptic efficacy values were the product *x*(*t*)*u*(*t*), where *x*(*t*) and *u*(*t*) are the time varying values representing the amount of neurotransmitter available, and the calcium concentration, respectively, as described above.

We measured the classification accuracy using cross-validation, in which we randomly selected 75% of trials for training the decoder and the remaining 25% for testing the decoder. For each of the eight motion directions, we randomly sampled, with replacement, 25 trials to train the decoder (from the 75% of trials set aside for training), and 25 trials to test the decoder (from the 25% of trials set aside for testing). From the 200 trials in the test set (25 time 8 directions), the accuracy was the fraction of trials in which the predicted motion direction matched the actual motion direction.

We used a bootstrap approach to determine statistical significance. We did so by repeated this sampling procedure 100 times to create a decoder accuracy distribution for each time point. The difference was deemed significantly greater than chance if 98 values were greater than chance (equivalent to P < 0.05 for a two-sided test).

For each network, we calculated decoding accuracies using a batch of 1024 trials in which the test motion directions were randomly sampled independently of the sample motion direction. This was in contrast to how we trained the network and measured behavioral accuracy, in which there was always a 50% chance that a test stimulus would match the sample. We note that the pattern of neural and synaptic activity generated by a sample stimulus will be similar to the pattern generated by a matching test stimulus. Thus, if matching test stimuli occur more frequently than chance, our sample decoding accuracy during and after the test stimuli would be artificially elevated.

### Measuring the contribution of neuronal activity and synaptic plasticity towards solving the task

To measure how network models used information maintained in neuronal activity and in dynamic synaptic efficacies to solve the task, we used a shuffling procedure as follows. At the time point right before test onset (or right before the third test onset for the A-B-C-A/A-B-B-A tasks), we either 1) did not shuffle any activity, 2) shuffled the neuronal activity between trials, or 3) shuffled the synaptic efficacies between trials. We shuffled between trials as opposed to between neurons because neurons can have different baseline activity levels, and shuffling this activity can significantly perturb the network and degrade performance, even if no information is maintained in their activity. We then simulated the network activity for the remainder of the trial using the saved input activity, and measured the performance accuracy by comparing the activity of the three output neurons to the target output activity during the test period. We performed this random shuffling 100 times, and measured the mean performance accuracy for all three conditions. The rationale behind this analysis is that if the network was exclusively using information maintained in neuronal activity to solve the task, then shuffling neuronal activity between trials should devastate performance, while shuffling synaptic efficacies should have little effect. Similarly, if the network was exclusively using information maintained in synaptic efficacies to solve the task, then shuffling synaptic efficacies between trials should devastate performance, while shuffling neuronal activity should have little effect. If the network was using information maintained in both neuronal activity and synaptic efficacies, then shuffling either should lead to significant decreases in performance.

### Tuning similarity index

We measured the similarity between sample and test stimuli encoding in the A-B-C-A and A-B-B-A tasks (Figure 5) using a similarity index we previously employed to study the relation between functional clustering and mnemonic encoding^10^. To calculate this index, we first modelled the synaptic efficacy for all neurons as a linear function of the sample or test motion direction, represented by the unit vector *d*:

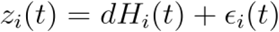

where *∊_i_*(*t*) is a Gaussian noise term and the vector *H_i_*(*t*) relates the stimulus direction to the synaptic efficacy.

The angle of *H_i_*(*t*) is the preferred direction of the neuron at time t, and its magnitude indicates the change in synaptic efficacy from baseline when the stimulus matches the preferred direction of the synapse, Thus, the preferred direction of a synapse, represented as a unit vector, is

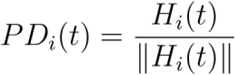

We can calculate how well this linear model fit the data for each synapse *i* and time point *t*, indicated by *w_i_*(*t*), by comparing the variance in the residuals with the variance in the synaptic efficacy:

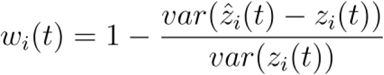

where the fitted synaptic efficacy is determined by the linear model: 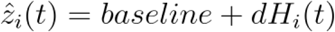.

For each synapses, we calculated these values for both the sample and first test motion direction, and then calculated the tuning similarity as the dot-product between their preferred sample and test motion directions directions, weighted by the geometric mean of their normalized linear model fits:

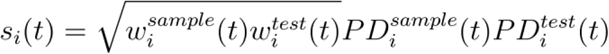

Finally, we calculated the similarity tuning index as the sum of the similarity scores for all synapses, divided by the sum of the geometric means of their respective linear model fits:

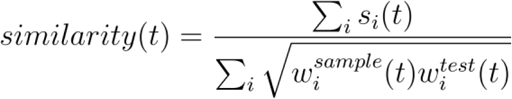

For the analysis examining whether preferred sample motion directions rotate during the DMRS task (Figure S2), we calculated how preferred sample motion directions changed across time based on the calculations above. To do so, we first represented the selectivity and preferred sample direction of neuronal activity or synaptic efficacies in complex form:

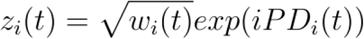

We then calculated the weighted change in preferred sample directions between synapses at different times:

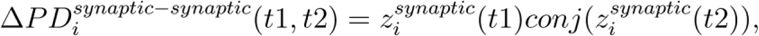
or the weighted change in preferred sample directions between neurons and their associated synapses at different times:

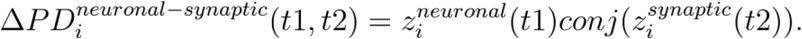

We then calculated the weighted histogram by binning the angular differences (*angle*(*ΔPD*)) weighted by their absolute value *(abs*(*ΔPD*)).

When calculating the tuning similarity index for Figure 7, we note that the delay period and the number of test stimuli differed between tasks. Thus, to calculate the metric in a standard way across all tasks, we tested all networks with a sample stimulus lasting 500 ms followed by a 1000 ms delay.

### Category tuning index

The category tuning index (CTI) measured the difference in synaptic efficacy (averaged across all trials for each direction) for each neuron between pairs of directions in different categories (a between category difference or BCD) and the difference in activity between pairs of directions in the same category (a within category difference or WCD)^51^. The CTI was defined as the difference between BCD and WCD divided by their sum. Values of the index could vary from +1 (which indicates strong binary-like differences in activity to directions in the two categories) to −1 (which indicates large activity differences between directions in the same category, no difference between categories). A CTI value of 0 indicates the same difference in firing rate between and within categories.

### Statistics

We trained 20 networks for each task in order to assess the variability between different network solutions. All networks were initialized with different sets of random weights. We report mean ± standard deviation throughout the paper unless otherwise noted. We measured correlation using the Pearson correlation coefficient, except for Figure 7, where we used the Spearman correlation coefficient because of the small sample size (N = 10). The data distribution was assumed to be normal, but this was not formally tested. No data points were excluded from the analyses, and no statistical methods were used to predetermine sample sizes, since all networks were able to solve the task with > 90% accuracy. Data collection and analysis were not performed blind to the conditions of the experiments, as this did not apply to our simulations. Data collection and assignment to experimental groups also did not apply, since all networks were equivalent before training.Table 1 Parameters used for network architecture and training.

**Table 1.** Parameters used for network architecture and training

| Parameter | Symbol | Value |
| --- | --- | --- |
| Learning rate | $\nu$ | 0.02 |
| Neuron time constant | $\tau$ | 100 ms |
| Time step (training and testing) | $\Delta t$ | 10 ms |
| Standard deviation of input noise | $\sigma_{in}$ | 0.1 |
| Standard deviation of recurrent noise | $\sigma_{rec}$ | 0.5 |
| L2 penalty term on firing rates | $\beta$ | 0.02 (0.005 for dual task) |
| STP neurotransmitter time constant | $\tau_x$ | 200 ms / 1500 ms<br>(facilitating / depressing) |
| STP calcium time constant | $\tau_u$ | 1500 ms / 200 ms (facilitating /<br>depressing) |
| STP calcium increment | $U$ | 0.15 / 0.45<br>(facilitating / depressing) |
| Number of neurons (input layer, recurrent layer, output layer) | $N_{in}, N_{rec}, N_{out}$ | 36, 100, 3 |
| Gradient batch size | $N_{trials}$ | 1024 |
| Number of batches during training |  | 2000 (4000 for dual task) |

## Supplementary Information

### Synaptic decoding accuracy for the A-B-C-A task

As noted in the Results, the synaptic decoding accuracy for the A-B-C-A task was relatively lower compared to other tasks, which was because of how we calculated decoding accuracy (see Methods). When training our networks on the A-B-C-A task, matching test stimuli, which can boost the sample information maintained in short-term memory, occur at least once during a trial with 87.5% probability 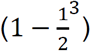. However, when calculating decoding accuracy, we randomly sampled test motion direction independently of sample directions, so that matching test stimuli occur at least once during a trial with 30.0% probability 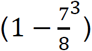. We sampled independently when calculating decoding accuracy because a sample stimulus and a matching test stimulus generate similar patterns of neural and synaptic activity, meaning that sample decoding accuracy would be artificially high if matching test stimuli occurred more frequently than chance. Thus, synaptic decoding accuracy in A-B-C-A task likely appears lower than expected because networks receive less matching test stimuli than they receive during training.

### Effect of noise and regularization

In the brain, neurons receive inputs from a wide range of cortical sources, from both local and distant areas, as well from a variety of subcortical structures. To model the impact of these diverse inputs, in which many might be unrelated to the task at hand, we added Gaussian noise directly to the RNN neurons, and to the input neurons (see Methods). We noticed that increasing the amount of noise decreases the reliability of decoding motion direction from neuronal activity, which encourages RNNs to rely more heavily upon information maintained in synaptic efficacies to solve the task. Additionally, we added a penalty term on the mean squared neuronal activity values of the recurrent neurons to encourage networks to solve the task using sparse neuronal activity. Increasing this penalty term also encourages RNNs to rely more heavily upon information maintained in synaptic efficacies, and less upon neuronal activity, to solve the task.

We obviously do not have access to effective noise levels in the brain, nor on how much pressure they are under to minimize the metabolic cost associated with generating action potentials. However, it is worth noting that how information maintained in short term memory is distributed between persistent activity compared to synaptic efficacies depends on these values, which might vary between species, cortical areas, etc.

**Figure S1.**
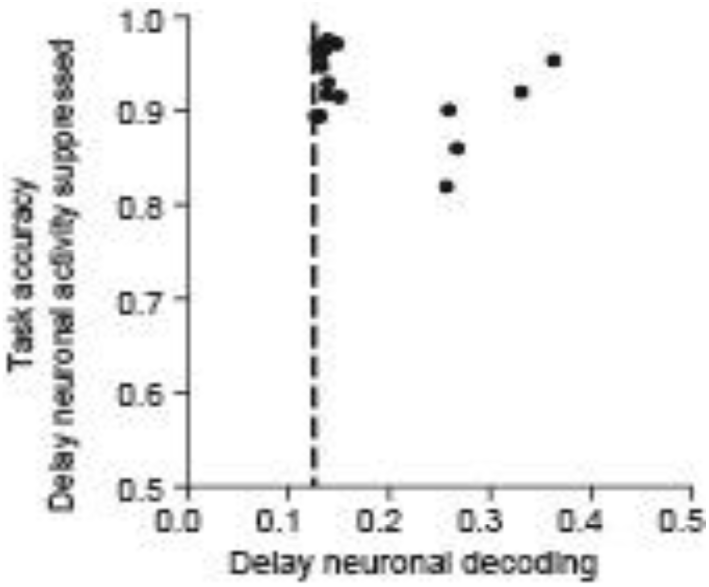
To confirm that most networks maintained information solely in synaptic efficacies during the DMS task, we measured task performance on trials during which we set neuronal activity for all recurrently connected neurons to zero for the last 100 ms of the delay period. Scatter plot shows the mean neuronal decoding accuracy during the last 100 ms of the delay period (x-axis) against the behavioral accuracy after suppressing all neuronal activity during the last 100 ms of the delay period (y-axis) for all networks that solved the DMS task. Each circle represents the results from one of the 20 networks. Suppressing neuronal activity during the end of the delay period had little effect on many networks, with accuracy remaining >90% for 16 out of 20 networks. Thus, the sample direction could be silently maintained in the synaptic efficacies across the delay period for most networks, which could then be read out when the network was driven by upcoming test stimulus.

**Figure S2.**
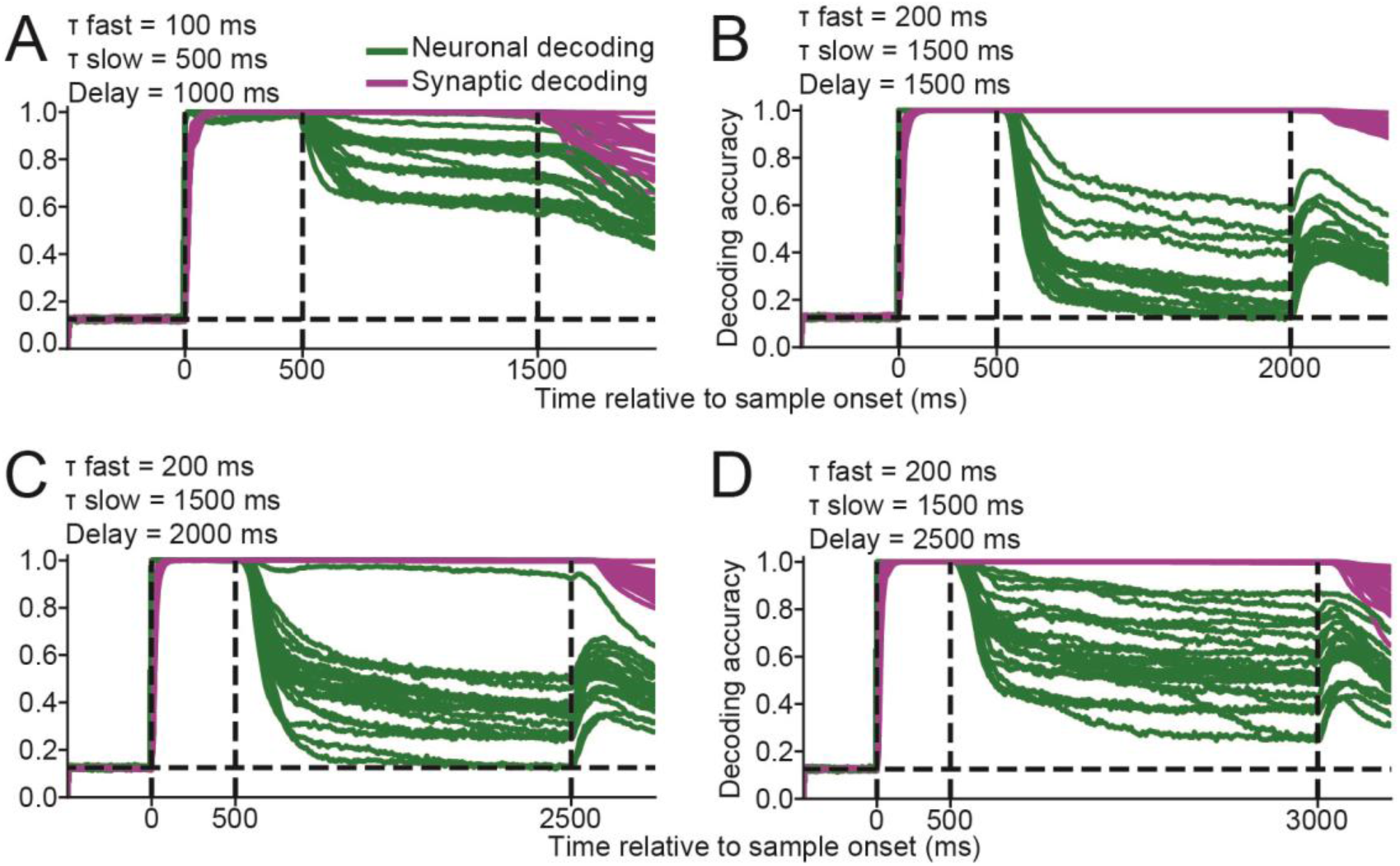
Exploring the conditions under which activity-silent encoding exists. **(A)** The time course of the sample direction decoding accuracy, calculated using neuronal activity (green curves) or synaptic efficacy (magenta curves) are shown for all twenty networks that successfully solved the DMS task with the STP time constants of 100 and 500 ms. Neuronal decoding accuracy was significantly greater than chance (P>0.05, bootstrap) during the last 100 ms of the delay period for all 20 networks. **(B-D)** Same as (A), with STP time constants set to 200 and 1500 ms, but with a delay period of 1500 (B), 2000 (C), and 2500 ms (D), respectively. Neuronal decoding accuracy was not significantly greater than chance (P>0.05, bootstrap) during the last 100 ms of the delay period for 8, 2, and 0 networks out of 20. Thus, networks did not solve the DMS task with a delay periods of 2500 ms using activity-silent memory.

**Figure S3.**
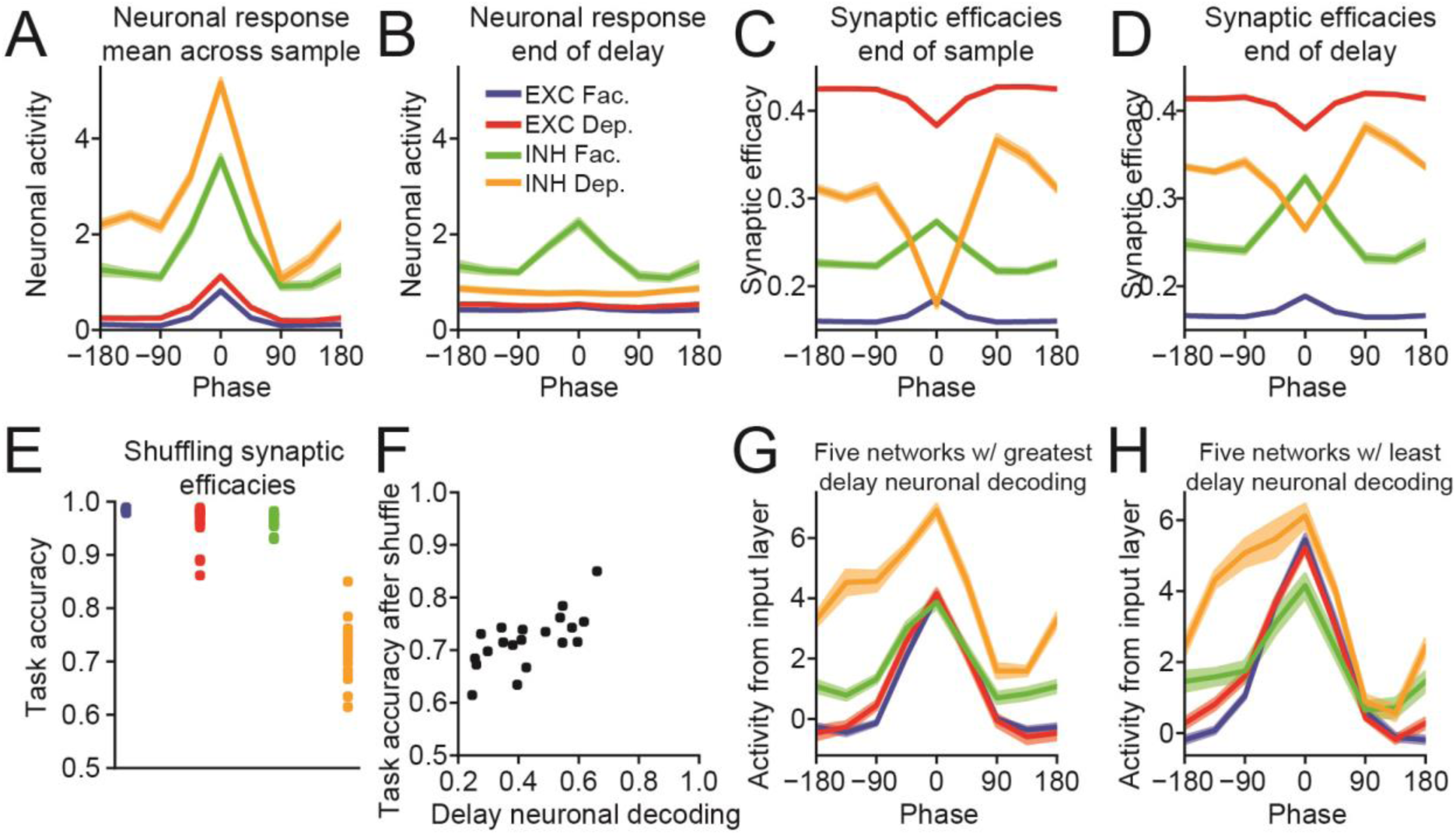
Understanding how networks solve the DMRS task with a 90° counterclockwise rotation. Same as Figure 3, except that networks were trained to solve a DMRS task in which the matching test stimulus was rotated 90° counterclockwise from the sample direction. **(A)** The neuronal sample tuning curves were calculated for four groups of neurons (excitatory neurons with facilitating synapses, blue; excitatory neurons with depressing synapses, red; inhibitory neurons with facilitating synapses, green; inhibitory neurons with depressing synapses, orange) from all 20 networks that solved the task. Neuronal activity was averaged across the entire sample period, and the tuning curves were centered around the sample direction that generated the maximum response (i.e. the preferred direction). Error bars (which are small and difficult to see) indicate one SEM. **(B)** Same as (A), except that neuronal activity at the end of the delay period was used to calculate the tuning curves. In order to compare how neural activity evoked during the sample evolves across the delay period, tuning curves were aligned to the preferred directions calculated in (A). **(C)** Same as (B), except that synaptic efficacies at the end of the sample period were used to calculate the tuning curves. As above, the preferred directions are the same as those used in (A). **(D)** Same as (C), except that synaptic efficacies at the end of the delay period were used. **(E)** Task accuracy after shuffling synaptic efficacies at the end of the sample period for each of the four groups of neurons. Each dot represents the accuracy from one network. **(F)** Scatter plot showing the neuronal decoding accuracy measured at the end of the delay period (x-axis) against the task accuracy after shuffling the synaptic efficacies of inhibitory neurons with depressing synapses at the end of the sample period (y-axis). Each dot represents the results of one of the 20 networks. **(G)** Tuning curves showing the activity received from the neurons in the input layer, which was calculated by multiplying the motion direction tuning of the neurons in the input layer with the input-to-recurrent weight matrix. Results were averaged across the 5 networks with the greatest neuronal decoding accuracy during the last 100 ms of the delay period. The four curves thus indicate the mean amount of input each group of recurrent neurons (excitatory or inhibitory, facilitating or depressing) receives from the input layer for each direction. As above, the preferred directions are the same as those used in (D). (H) Same as (G), except showing the input tuning curves averaged across the e 5 networks with the lowest neuronal decoding accuracy during the last 100 ms of the delay period.

**Figure S4.**
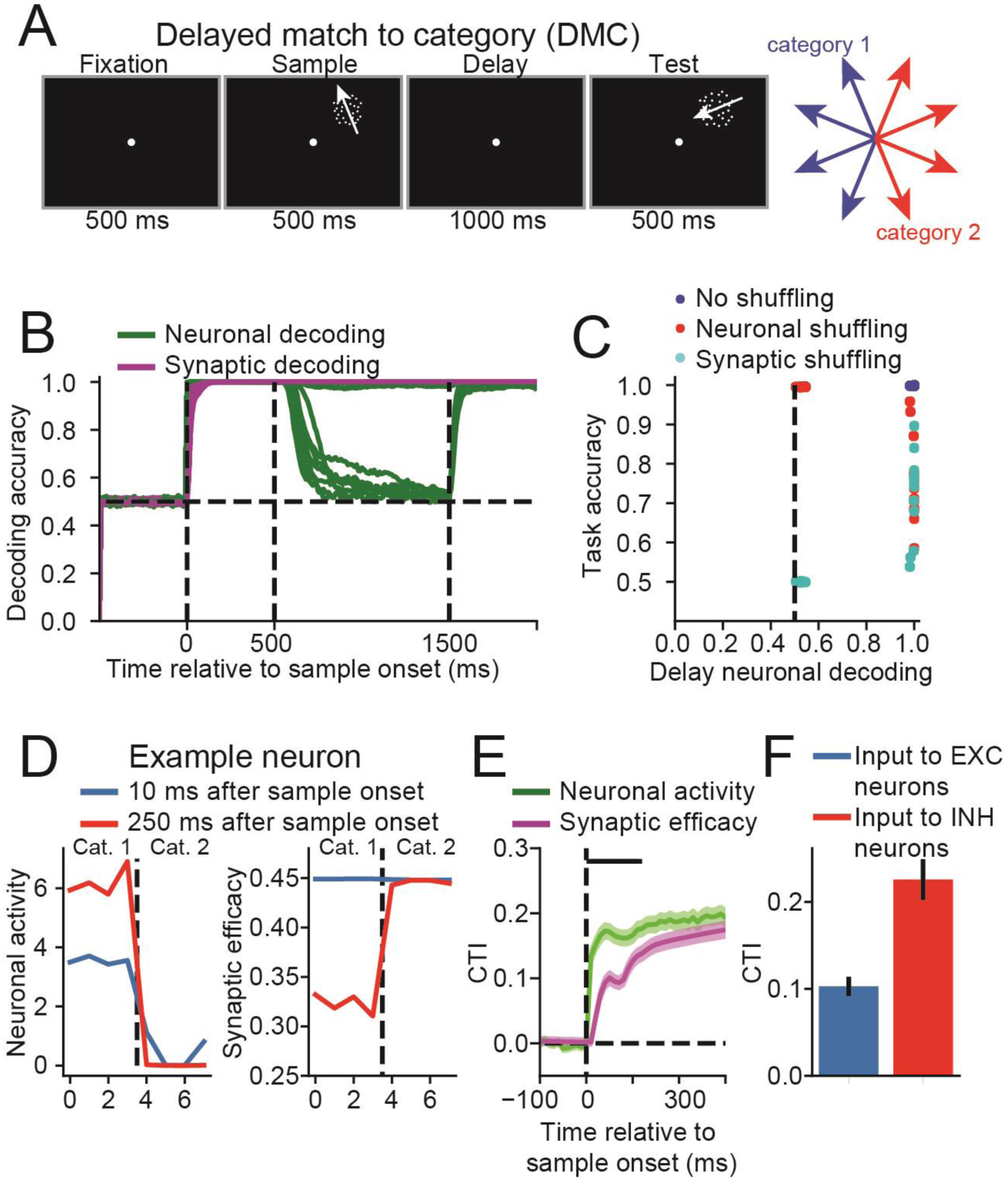
Neuronal and synaptic category selectivity during the delayed match-to-category task. **(A)** The delayed match-to-category task is similar to the DMS task, except that the network was trained to indicate whether the category membership of the sample and test stimuli matched. The eight motion directions were divided into two categories, indicated by the red and blue motion directions. **(B)** The time course of the sample direction decoding accuracy, calculated using neuronal activity (green curves) or synaptic efficacy (magenta curves) are shown for all twenty networks that successfully solved the DMS task. The dashed vertical lines, from left to right, indicate the sample onset, offset, and end of the delay period. The neuronal decoding accuracy, measuring during the last 100 ms of the delay period, were not significantly greater than chance for 7 out of 20 networks. **(C)** Scatter plot showing the neuronal decoding accuracy measured at the end of the delay (x-axis) versus the behavioral accuracy (y-axis) for all 20 networks trained on this task (blue circles), the behavioral accuracy for the same 20 networks after neuronal activity was shuffled right before test onset (red circles) or synaptic efficacies were shuffled right before test onset (cyan circles). Thus, for each blue circle, there is a corresponding red and cyan circle with matching neuronal decoding accuracy. The dashed vertical line indicates chance level decoding accuracy. (D) Motion direction tuning of a single example neuron showing the mean neuronal activity (left panel) or mean synaptic efficacy (right panel) as a function of the motion direction index (x-axis). The dashed vertical line divides the motion directions into two categories. The blue curves show the tuning function calculated 10 ms after sample onset, and the red curves shows the tuning function calculated 250 ms after sample onset. This neuronal activity for this example neuron is category-selective 10 ms after sample onset, showing high, mostly uniform, activity for category 1 and uniformly weaker activity for category 2. The synaptic efficacies for this example neuron were not category selective 10 ms after sample onset. (E) The time course of the mean category-tuning index (CTI, see Methods) using neuronal activity (green curve) and synaptic efficacy (magenta curve) averaged across all 7 networks whose neuronal decoding accuracy during the last 100 ms of the delay period were not significantly greater than chance. (F) The mean CTI of the activity projected from the motion direction tuned neurons in the input layer into excitatory (blue) and inhibitory neurons (red). Both values are significantly above zero (P < 10^−6^ for both, two-sided, t-test), indicating that connection weights from the input neurons are at least partially responsible for the category selectivity observed in (E).

**Figure S5.**
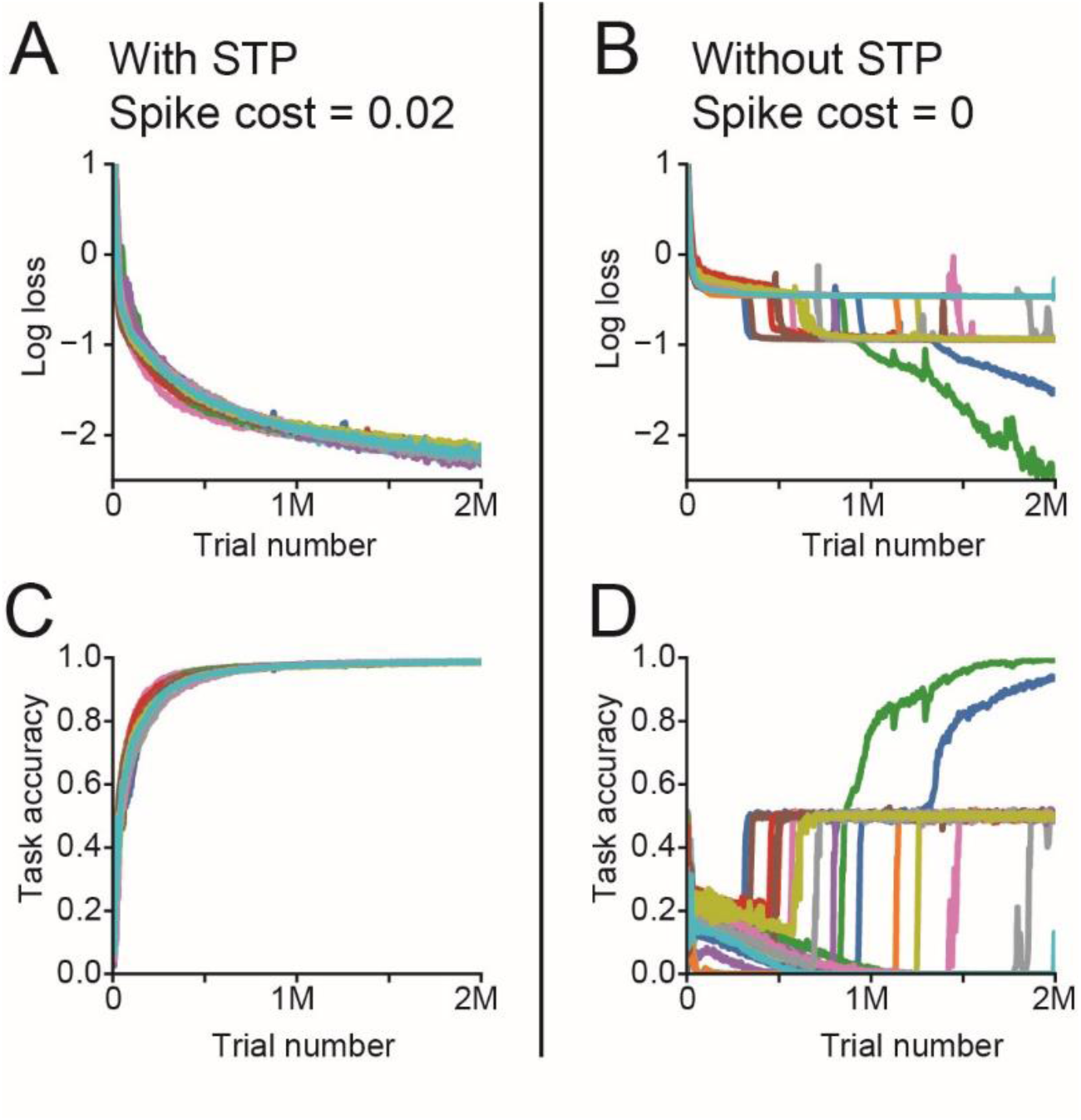
The evolution of the loss value and the task accuracy during the training of networks with and without STP on the DMS task. **(A)** The time course of the logarithm of the loss value during training of the networks with STP. These 20 networks were the same used for Figure 2, and each trace indicates the loss value of one of the 20 networks. All curves were smoothed with a boxcar filter of size 5120 trials (5 training batches). As expected, the loss value decreases throughout training. **(B)** Similar to (A), except for networks without STP and with spike cost set to zero. The loss value initially decreases at the start of training, but then stalls for many networks. **(C)** The time course of the task accuracy the training of networks with STP. All 20 networks successfully learned to perform the task with accuracy >95% after 1,000 training batches (~1,000,000 trials). **(D)** Similar to (C), except for networks without STP and with spike cost set to zero. Only 2 out of the 20 networks successfully learned to perform the task with accuracy >90% after 2,000 training batches (~2,000,000 trials).

